# The PLETHORA/PIN-FORMED/AUXIN network mediates terminal prehaustorium formation in the parasitic plant *Striga hermonthica*

**DOI:** 10.1101/2021.06.24.449531

**Authors:** Ting Ting Xiao, Gwendolyn K. Kirschner, Boubacar A. Kountche, Muhammad Jamil, Savina Maria, Vinicius Lube, Victoria Mironova, Salim al Babili, Ikram Blilou

**Affiliations:** BESE Division, Plant Cell and Developmental Biology, King Abdullah University of Science and Technology, Thuwal, Kingdom of Saudi Arabia; BESE Division, The BioActives Lab, King Abdullah University of Science and Technology, Thuwal, Kingdom of Saudi Arabia; Institute of Cytology and Genetics, Lavrentyeva Avenue 10, Novosibirsk, 630090, Russian Federation; Novosibirsk State University, 2 Pirogova Street, Novosibirsk, 630090, Russian Federation

**Keywords:** *Striga hermonthica*, root differentiation, auxin, PIN proteins, mathematical model, root meristem, haustorium

## Abstract

The parasitic plant *Striga hermonthica* invades the host root through the formation of a haustorium and has detrimental impacts on cereal crops. The haustorium is derived directly from the differentiation of the *Striga* radicle. Currently, how *Striga* root cell lineages are patterned and the molecular mechanisms leading to radicle differentiation shortly after germination remain unclear. In this study, we determined the developmental-morphodynamic programs that regulate terminal haustorium formation in *S. hermonthica* at spatiotemporal and cellular resolutions. We showed that in *S. hermonthica* roots, meristematic cells first undergo multiplanar divisions, which decrease during growth and correlate with reduced expression of the stem cell regulator *PLETHORA1.* We also found that PIN-FORMED (PIN) proteins undergo a shift in polarity. Using the layout of the root structure and the polarity of outer-membrane PIN proteins, we constructed a mathematical model of auxin transport that explains the auxin distribution patterns observed during *S. hermonthica* root growth. Our results reveal a fundamental molecular and cellular framework governing the switch of *S. hermonthica* roots from the vegetative to the invasive state by inducing meristem differentiation through auxin excretion to the environment and explain how asymmetric PIN polarity controls auxin distribution to maintain meristem activity and sustain root growth.

## Introduction

The parasitic plant *Striga hermonthica,* commonly known as purple witchweed, is an obligate root hemiparasite belonging to the Orobanchaceae family that causes massive yield losses in agronomically important cereal crops such as sorghum, maize, and millet. It is considered a major threat to global food security (Pennisi, 2015), particularly in sub-Saharan Africa (Ejeta, 2007; Kountche et al., 2016). *Striga* seeds can only germinate in close proximity to the host roots as it depends on stimulants, mainly strigolactones, that are produced by the host roots (Cook et al., 1972; Al-Babili and Bouwmeester, 2015). *Striga* sp. invades its host by forming a haustorium, a hairy structure that is central to successful invasion. Members of the Orobanchaceae family employ two types of haustoria: a primary or terminal haustorium that develops from the differentiation of the root meristem shortly after germination and a lateral haustorium that develops from the de-differentiation of cells from the cortex, similar to *de novo* organ formation, such as in lateral roots or root nodules (Joel and Losner-Goshen, 1994; Goyet et al., 2019).

The initiation of prehaustorium formation requires host-derived molecules from plant root exudates known as haustorium-inducing factors (HIFs). One of the active compounds is quinone 2,6-dimethoxy-1,4-benzoquinone (Chang and Lynn, 1986), which is recognized by leucine-rich repeat receptor-like kinases in the hemiparasite *Phtheirospermum japonicum* (Laohavisit et al., 2020). The prehaustorium invades the host by applying mechanical pressure to the host cells in combination with enzymatic secretion, thereby allowing the prehaustorium to enter the intercellular space and colonize the host root (Perez-de-Luque, 2013). Upon successful invasion, a haustorium is formed, through which the parasite establishes a tight connection with the host’s vascular system to acquire water and nutrients, enabling it to grow and proliferate at the expense of the host (Yoshida et al., 2016). The auxin biosynthesis gene *YUCCA3* has been found to be essential for lateral haustorium formation in *P. japonicum* (Ishida et al., 2016), and the establishment of the xylem bridge that connects the parasite to the host is mediated by polar auxin transport; the PIN-FORMED (PIN) proteins, which are efflux carriers, play a major role in this process (Wakatake et al., 2020).

The process of lateral haustorium formation in the facultative parasite *P. japonicum* has been extensively studied, and the mechanisms underlying its development are slowly emerging owing to the availability of genetic tools and cell type-specific reporters. The lack of these tools in other parasitic species hampers advancement in the understanding of the developmental programs underlying the formation of the lateral and terminal haustoria. Nevertheless, comprehensive transcriptome analysis of *Striga* roots from germination to haustorium formation has revealed the differential expression of developmental genes at the early stages, followed by an increase in the expression level of genes involved in auxin transport and response (Yoshida et al., 2019). Despite these recent advances, there is a lack of comprehensive analysis of the cellular and molecular morphodynamics *of Striga* roots that lead to terminal haustorium formation.

In this study, we describe the development of the terminal haustorium in *S. hermonthica* and provide new insights into the mechanisms underlying its morphodynamics during growth. We showed that after germination, *S. hermonthica* develops a root meristem with an anatomy that resembles that of the model plant *Arabidopsis thaliana* (Hood et al., 1998; Raghavan and Okonkwo, 1982). We found that meristematic cells undergo multiplanar cell division, including those at the quiescent center (QC) position within a stem cell niche marked by the stem cell regulator *PLETHORA1 (PLT1).* Prior to haustorium formation, the rate of meristematic cell division decreases, the cells elongate, and root hairlike structures emerge from the root tip. We demonstrated that terminal haustorium formation is correlated with basal epidermal polarity of the auxin transporters PIN1/PIN2. Based on the root structure layout of *S. hermonthica* and the observed basal PIN polarity, we constructed a mathematical model that depicted the auxin distribution pattern in *S. hermonthica* roots and recapitulated the expected decrease in auxin levels at the elongated stage resulting from basal PIN polarity, followed by auxin depletion prior to the haustorium stage. Consistent with these results, *S. hermonthica* induced an increase in the auxin response when it was in contact with Arabidopsis roots. These results establish a mechanistic model that explains the switch of the *S. hermonthica* root from the vegetative to the invasive state.

## Material & Methods

### S. hermonthica seed germination and growth conditions

*S. hermonthica* seeds were collected from sorghum fields in Sudan. The seeds were first surface-sterilized with 50% commercial bleach solution (NaOCl) for 12 min on a rotary shaker. The seeds were then washed six times with double-autoclaved water and placed on nylon filter paper moisturized with sterile water. Preconditioning was performed at 30°C in the dark for 10 days in sterile water. Germination was induced by application of 2.5 μM roc-GR24, either directly on filter paper or into ½ Murashige and Skoog (MS) growth medium.

### Measurement of meristem size

Meristem length was measured from the cell above the root cap to the first elongated cell within the epidermis. All box plots were generated using IBM SPSS Statistics for Windows, version V26.

### Evaluation of cell division using EdU staining

EdU was added at a final concentration of 5 μM to seeds treated with GR24 and incubated in MS plant growth medium supplemented with 1% sucrose, for 90 min (Fig. 2) or 12 h (Fig. 3). EdU staining was performed using the Click-iT EdU Alexa Fluor 488 Imaging Kit (C10637; Invitrogen, ThermoFisher scientific, USA)) as described previously (Cruz-Ramírez et al., 2013; Kirschner et al., 2017). The samples were cleared in chloral hydrate solution for several days (Truernit et al., 2008), followed by cell wall and DNA counterstaining using mPS-PI and Hoechst solution (5 μg/ml; Invitrogen) or SCRI Renaissance staining SR2200, as described previously (Musielak et al., 2015). Images were captured using an inverted confocal microscope (Leica SP8 or LSM 880; Carl Zeiss).

### Quantification of cell division rate

The standard sketches of *S. hermonthica* roots were generated based on confocal images. EdU-stained images were classified into different groups based on their developmental stages. Then, the images were overlaid on the standard sketch of their corresponding stages. Subsequently, the dividing cells were mapped onto the cells on the sketches. Finally, heatmaps were generated based on the division frequency of the cells in the standard sketches.

### Starch staining using Lugol’s iodine

Seedlings were incubated with Lugol’s iodine and mounted in chloral hydrate solution as described previously (Van den berg et al., 1997). Images were acquired using a transmitted light illumination Leica DM2500 microscope with an HC PL FLUOTAR 40x objective.

### Staining of cell walls and nuclei

To visualize the cell wall, SCRI Renaissance staining SR2200 was used as described previously (Musielak et al., 2015). For visualization of lignin and suberin, *S. hermonthica* roots were embedded in low-melting-point agar and sectioned using a vibratome as described previously (Xiao et al., 2019). The cells were stained with basic fuchsine or berberine hemisulfate as described previously (Ursache et al., 2018).

Sections were imaged using an inverted confocal microscope (Leica SP8 or LSM 880; Carl Zeiss) with 405 nm excitation for SCRI and 469 nm emission and 488 nm excitation for berberine and 546 nm emission. For basic fuchsine, 561 nm excitation and 617 nm emission were used.

### Preparation of root extract

*Arabidopsis* and rice seeds were sterilized, germinated, and grown on ½ MS medium for 3 weeks. The roots were harvested in liquid nitrogen and ground. The resulting powder was dissolved in sterile water to a concentration of 5% (w/v) as described previously (Wada et al., 2019).

### RNA in situ hybridization

*S. hermonthica* homolog genes were identified by blasting the CDS *of Arabidopsis* genes at the Parasitic Plant Genome Project website (http://ppgp.huck.psu.edu) (Yang et al., 2015). The expressed sequence tags were also blasted to the newly released genome (Yoshida et al., 2019). The *S. hermonthica* sequences with the best hit were used for generating and cloning the probes. For RNA probes, cDNA fragments with lengths of 500-1,340 bp were amplified with Phusion polymerase using *S. hermonthica* cDNA obtained from seedlings. The obtained products were cloned into pGEM-T Easy Vector (Promega). The probes were synthesized by amplifying the cloned fragment with M13 primers. The purified PCR product was then used for an in vitro transcription reaction using the DIG RNA Labeling Mix (#11175025910, Roche, Germany) and T7 and Sp6 RNA polymerase to generate both sense and antisense probes. Whole-mount RNA *in situ* hybridizations were performed in seedlings embedded in low-melting-point agarose and sectioned as described previously (Hejátko et al., 2006). The samples were imaged using the transmitted light illumination Leica DM2500 microscope (Leica Microsystems). The sequences of the primers used to obtain the cDNA fragments are listed in Table S2.

### Immunolocalization

Whole-mount immunolocalization experiments using anti-PIN1 (Gälweiler et al., 1998) and anti-PIN2 (Müller et al., 1998) antibodies were performed as described previously (Blilou et al., 2005). To detect root cap cells with LM8 (Plant probes, PP-008), auxins with anti-IAA (Agrisera product no. AS06 193), and cytokinins with anti-cytokinin (Agrisera product no. AS09 437), immunolocalization was performed on paraffin sections as described previously (Forestan and Varotto, 2013). The antibodies were diluted as follows: 1/1000 dilution for anti-PIN1 and -PIN2 and 1/500 for auxins and cytokinins. For secondary antibodies, goat anti-rabbit IgG and Alexa Fluor 488 (ThermoFisher) at a dilution of 1/1000 were used. Images were acquired using a Leica Sp8 inverted confocal microscope.

### IAA treatments

*S. hermonthica* germination was induced with GR24 for 2 days. The seedlings were then transferred to media containing different concentrations of IAA of 10^-9^-10^-5^ M. The cell division rate in the meristem was monitored daily for 4 days after IAA treatment.

### Three-dimensional (3D) time-lapse imaging and guantification of fluorescence intensity

*Arabidopsis* seedlings containing *DR5::vYFP* and *DII::venus* were grown for 5 days and were put in contact with germinated *S. hermonthica* for 48 h until prehaustorium formation. To acquire 3D time-lapse images using *DR5::vYFP,* Z-stack images were recorded using Zen software at a rate of 50 frames with 104 slices and an average of 4 frames/s using a 10x plan Apochromat objective lens. To acquire fluorescence time-lapse images of *Arabidopsis* roots using *DR5::vYFP* and *DII::venus,* the fluorescence was visualized when roots were put in contact with *S. hermonthica* and the signal was recorded daily for 48 h for each root. YFP was excited at 514 nm, and emission was recorded at 575 nm. The fluorescence intensity was then quantified using ImageJ software as described previously (Long et al., 2017, 2018).

### Phylogenetic analysis and gene IDs

To construct the PIN and PLT phylogenetic tree, the *Arabidopsis* PIN1 and PLT1 amino acid sequences were used for a BLAST search for homolog sequences in the *Striga asiatica* genome release with a cutoff value of e-value <2e-05 for PIN and <7e-55 for PLT. *Arabidopsis* PLT sequences were obtained from arabidopsis.org. Rice PLTs were obtained from a previous study (Li and Xue, 2011). The sequences were aligned using Muscle in MEGA X (Kumar et al., 2018), and the evolutionary history was inferred using the Maximum Likelihood method and JTT matrix-based model integrated in the software with a bootstrap value of 100.

The following gene IDs are mentioned in this study: *ShPLT1* (StHeBC3_20835.1, GER38334), *ShHISTONE H4* (StHe2GB1_80290), *ShCYCLIN B1,3* (StHeBC3_3205), *ShPIN1-1* (StHe61GB1_24115, GER51912), and *ShPIN1-2* (StHe3G2B1_75771, GER32949).

## Results

### Cellular organization of the *Striga* root meristem

After germination, the *Striga* radicle contains a meristem that has a simple and organized structure (Hood et al., 1998; Raghavan and Okonkwo, 1982). As growth progresses, *Striga* roots undergo anatomical changes that lead to the formation of a prehaustorium prior to host attachment (Joel and Gressel, 2013; Yoshida et al., 2016). To understand the developmental program of *Striga* embryonic roots, we induced seed germination using GR24, a synthetic analog of strigolactones that stimulates *Striga* germination, and performed an initial analysis in the absence of the host and HIFs. First, we categorized the developmental stages of *Striga* embryonic root growth. These stages were designated based on the cellular layout, tissue organization, and meristematic activity as follows (Fig. **1A-G**): a young stage, in which the root emerges from the seed coat; at this stage, the *Striga* radicle has a structured root meristem, similar to that of the model plant *A. thaliana* (Fig. 1B;G;P); a pre-elongated stage, in which cells shootward of the meristem increase in length and contribute to increasing the length of the radicle (Fig. 1C;H); an elongated stage, with only a few meristematic cells and with root hairs initiating on the lateral side of the epidermis above the meristematic cells (Fig. 1D;I); and a differentiated stage without a meristem and with root hairs emerging from the tip of the radicle (Fig. **1E;J**). In the young stage, *Striga* roots have a simple and organized structure with regular cell files, which is consistent with the results of previous studies (Hood et al., 1998; Raghavan and Okonkwo, 1982). From outside to inside, we could distinguish the epidermis, ground tissue (cortex and endodermis), and vasculature based on cell lineage (Fig. 1F;G;P). Similar to that in *A. thaliana,* we observed QC-like cells at the base of the vasculature and ground tissue layers, and distal to it, we observed a cell layer that we named the columella as well as epidermal precursor cells (CEPSCs) (Fig. **1F;G**, Fig. **S1**). The CEPSCs were protected by a single-layered root cap that we marked using the LM8 antibody, an epitope specific for xylogalacturonan that stains cells that are in the process of loosening from other tissues, such as those located at the root cap (Willats et al., 2004) (Fig. **1K-K’**). In Arabidopsis and other species, the distal columella comprises the columella stem cells and differentiated columella layers, of which only the differentiated columella layers accumulate starch granules (Van den berg et al., 1997). However, in *S. hermonthica,* we found starch granules in the distal root cells and the meristem, as evidenced using Lugol’s iodine and pseudo-Schiff propidium iodide staining (mPSI-PI) staining (Fig. **1L**, Fig. **2A”-E”**, Fig. **SI**). In addition, unlike in other plant species in which root cap cells detach and are continuously replenished during root growth (Kumpf and Nowack, 2015), we detected neither additional root cap cell layers nor root cap cell sloughing in *S. hermonthica* during root growth (Fig. **S1;** Fig. **2A-J**).

**Fig. 1:**
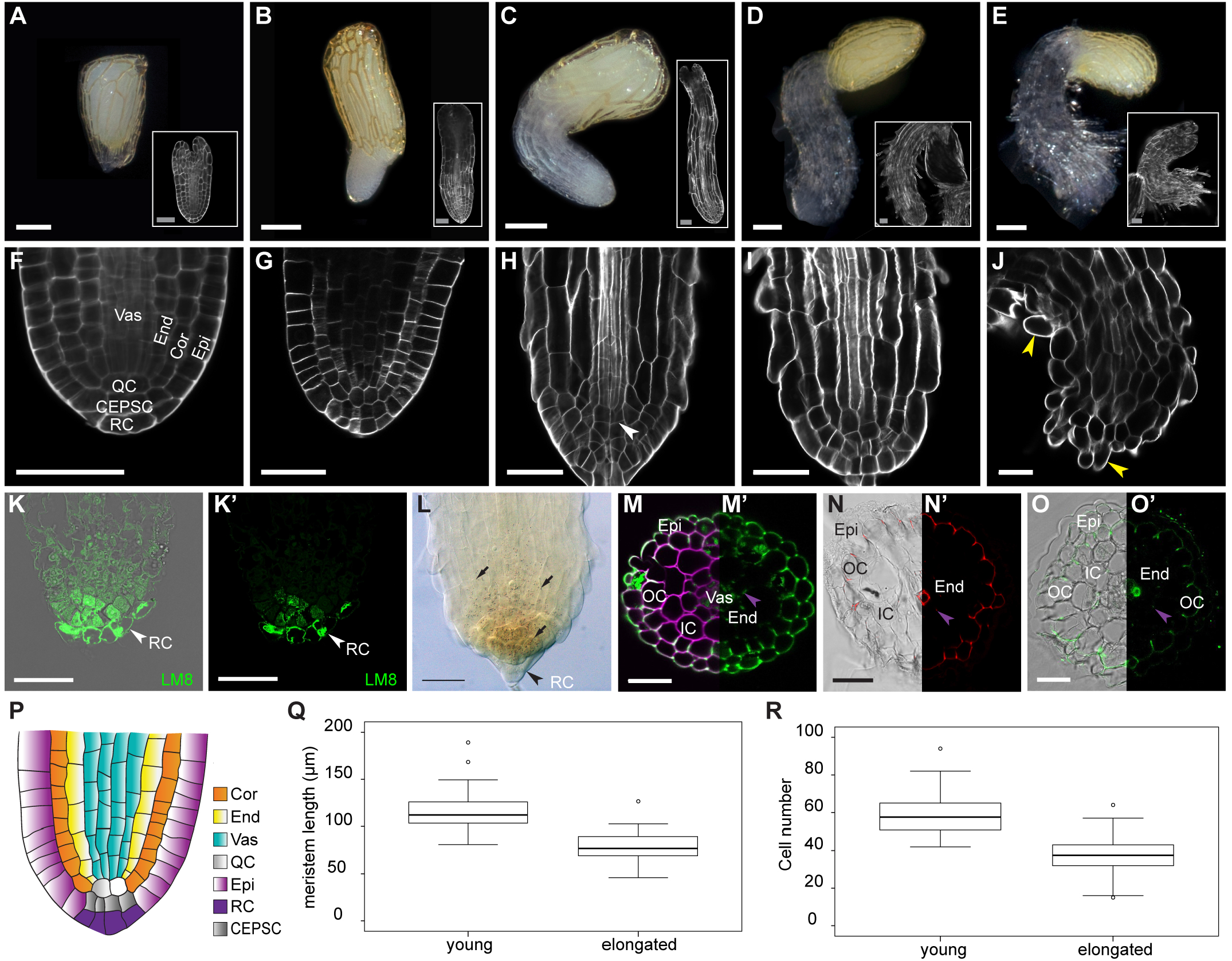
Cellular organization of the *Striga hermonthica* root meristem. A-E, Macrophotographs of *S. hermonthica* seedlings at the following different developmental stages: A, seed; B, young; C, pre-elongated; D, elongated; and E, differentiated. White rectangles show their respective confocal sections. F-J, Longitudinal confocal sections of the *S. hermonthica* meristem. K-K’, Immunolocalization using the LM8 antibody to mark the root cap in *S. hermonthica.* L, Lugol’s iodine stain showing stem cell differentiation. M-O’, Cross-sectional image of *S. hermonthica* with cell wall staining showing tissue composition. M, Renaissance stain. M’, Auto-fluorescence. N and N’, Basic fuchsine. 0 and O’, Berberine stain. P, Schematic representation of the *S. hermonthica* root meristem at the young stage. Q, R boxplots represent measurement of the meristem size (Q) and cell number (R) in young (number of plants, N = 86) and elongated (N = 55) *S. hermonthica* seedlings (24- and 48-h GR24 treatment). Horizontal lines indicate the medians. Box limits indicate the 25^th^ and 75^th^ percentiles. Whiskers extend to the 5^th^ and 95^th^ percentiles. T: B;C;G;H 12 h and D;E;I;J;M;N;O 48 h after GR24 treatment. Scale bars: *S. hermonthica* macrophotographs in (A-E): 200 μm; Images in (A-E; F-O’): 50 μm. Epi, epidermis; OC, outer cortex; IC, inner cortex, End, endodermis, Vas, vasculature; RC, root cap; Cor, cortex; QC, quiescent center; CEPSC, columella and epidermal precursor cells; arrows heads pointing to QC (H), root hairs(J), root cap(K-L) and casparian strips (M-O’).

**Fig. 2:**
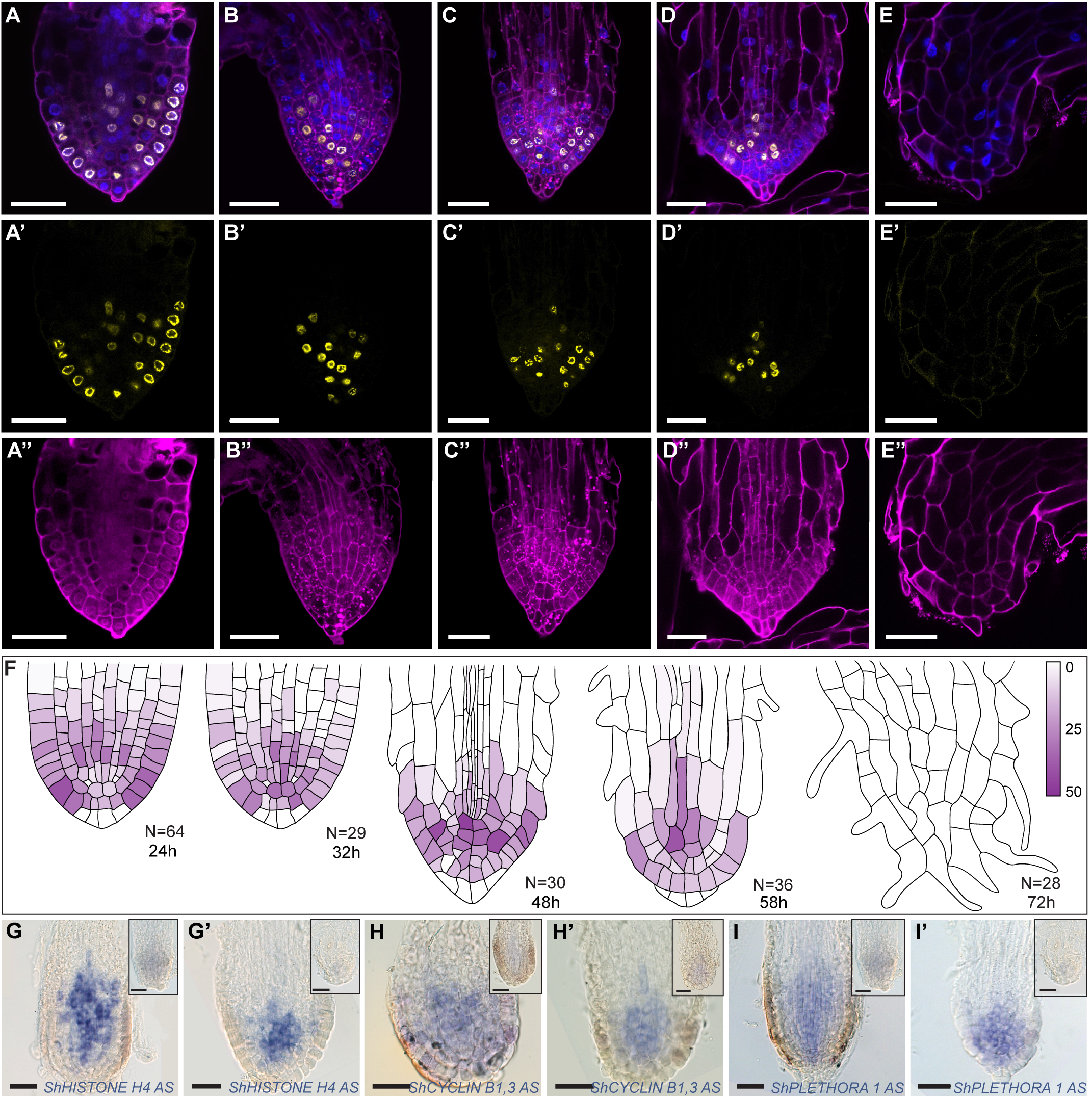
Haustorium formation correlates with meristem differentiation. A-E”, Confocal images of *Striga hermonthica* roots stained with the cell wall stain mPS-PI (A”-E”, purple), EdU (A’-E’; yellow), and the nuclear stain Hoechst (blue). A-B”, young stage; C-C”, pre-elongated stage, D-D”, elongated stage, E-E”, differentiated stage, F, quantification of the cell division rate during root differentiation, purple bar indicates the percentage of chance of that cell may undergo cell division (EdU marked nucleus /N*100). RNA *in situ* hybridization during meristem differentiation in longitudinal sections using *ShHistone-H4* (G-G’), *ShCYCLIN B1,3* (H-H’), and *ShPLETHORA1* (I–I’) probes. Roots in the upper right of the images represent the sense control for each probe. *S. hermonthica* seedlings with 24 or 48 h GR24 treatment. N = 30 for each probe. Scale bars: 50 μm.

During root growth, we observed anticlinal divisions within the meristem, particularly at the pre-elongated and elongated stages (Fig. S1). They occurred in the epidermis and at cells at the QC position, and periclinal divisions in the cortex and endodermis generated additional cell files (Fig. S1). In roots, tissue types can be defined by the composition of their cell wall (Ursache et al., 2018); therefore, we used different dyes to assess differences in cell wall composition in cross-sections of *S. hermonthica* roots. Using auto-fluorescence to detect the presence of lignin and suberin, we detected a bright fluorescent signal in the epidermis and adjacent layer, endodermis, and in a subset of vascular cells, suggesting the accumulation of lignin and suberin in these tissues (Fig. 1M;M’). These observations were confirmed using the fluorescent stains basic fuchsin and berberine to monitor lignin and suberin deposition, respectively (Fig. **1N-O’**).

### Meristem differentiation in *S. hermonthica*

To evaluate the growth dynamics at the different developmental stages, we first measured the size of the meristem and determined the number of cells in the young and elongated stages. We found that in the elongated stage, the meristem size was reduced and there were fewer cells in the meristematic zone compared with those in the young stage (Fig. **1Q;R**).

Next, we quantified the cell division rate during the different stages and in different tissue types using ethyl deoxyuridine (EdU), which is incorporated into cellular DNA during replication, marking cells in the S phase (Cruz-Ramírez et al., 2013) (Fig. **2A-E”**). We observed high meristematic cell division rates during the young stage (Fig. **2A-A’;F;** Fig. **S2).** At the pre-elongated and elongated stages, cells continued to divide at the distal meristem, including QC and columella stem cells (Fig. 2 **B-C’;F’).** However, prior to differentiation, we observed a rapid decrease in the cell division rate in the distal root apex (Fig. **2D;F**), followed by an arrest in meristematic activity (Fig. **2E-F;** Fig. **S2**). To verify these observations, we performed RNA *in situ* hybridization using *Striga HISTONE H4,* which is expressed during the progression of the G1-S phase of the cell cycle (Gutierrez, 2009; Fig. **2G;G’**), and the mitotic-specific G2-M phase *Striga CYCLIN-B1-3* (Bulankova et al., 2013; Fig. **2H;H’**). We found that the expression levels of both *ShHISTONE H4* and *ShCYCLIN-B1-3* were decreased as the *S. hermonthica* meristem underwent differentiation (Fig. **2G-H’**).

In *Arabidopsis,* the loss of activity of *PLT* leads to the loss of stem cell activity, the reduction of cell division rates, and differentiation of root meristem (Galinha et al., 2007). Therefore, we used *PLT* to evaluate whether progression to haustorium formation was correlated with the loss of stem cell activity. RNA *in situ* hybridization using a probe of the *Striga* homolog of *PLT1 (ShPLT1,* Fig. **S3**) revealed high mRNA levels in the meristem during progression to differentiation and a decrease in *ShPLT1* transcript levels during prehaustorium formation (Fig. **2I;I’**).

### Prehaustorium induction can occur in the absence of HIFs

Prehaustorium formation is triggered by host-derived HIFs. However, *Striga* roots can also grow in the absence of HIFs. To assess whether HIFs affect *Striga* root growth as well as their anatomy and tissue composition, we germinated *S. hermonthica* with GR24 and monitored the root growth and meristem cellular layout using plants grown in water or in the presence of non-host *(Arabidopsis)* and host (rice) extracts (Fig. **S4, S5).** We did not observe any changes in the growth dynamics between the treatments (Fig. **S4, S5).** Next, we assessed meristem activity using EdU staining and by monitoring the expression level *of ShPLT1* (Fig. **3A-K’**). We observed an increase in cell division rates and aberrant division plans in samples treated with root extracts, and the cells became elongated and larger, leading to the formation of thicker roots (Fig. **3B;C;E;F;** Fig. **S4).** We did not detect changes in the *ShPLT1* expression level between the treatments (Fig. **3J-K’**).

**Fig. 3:**
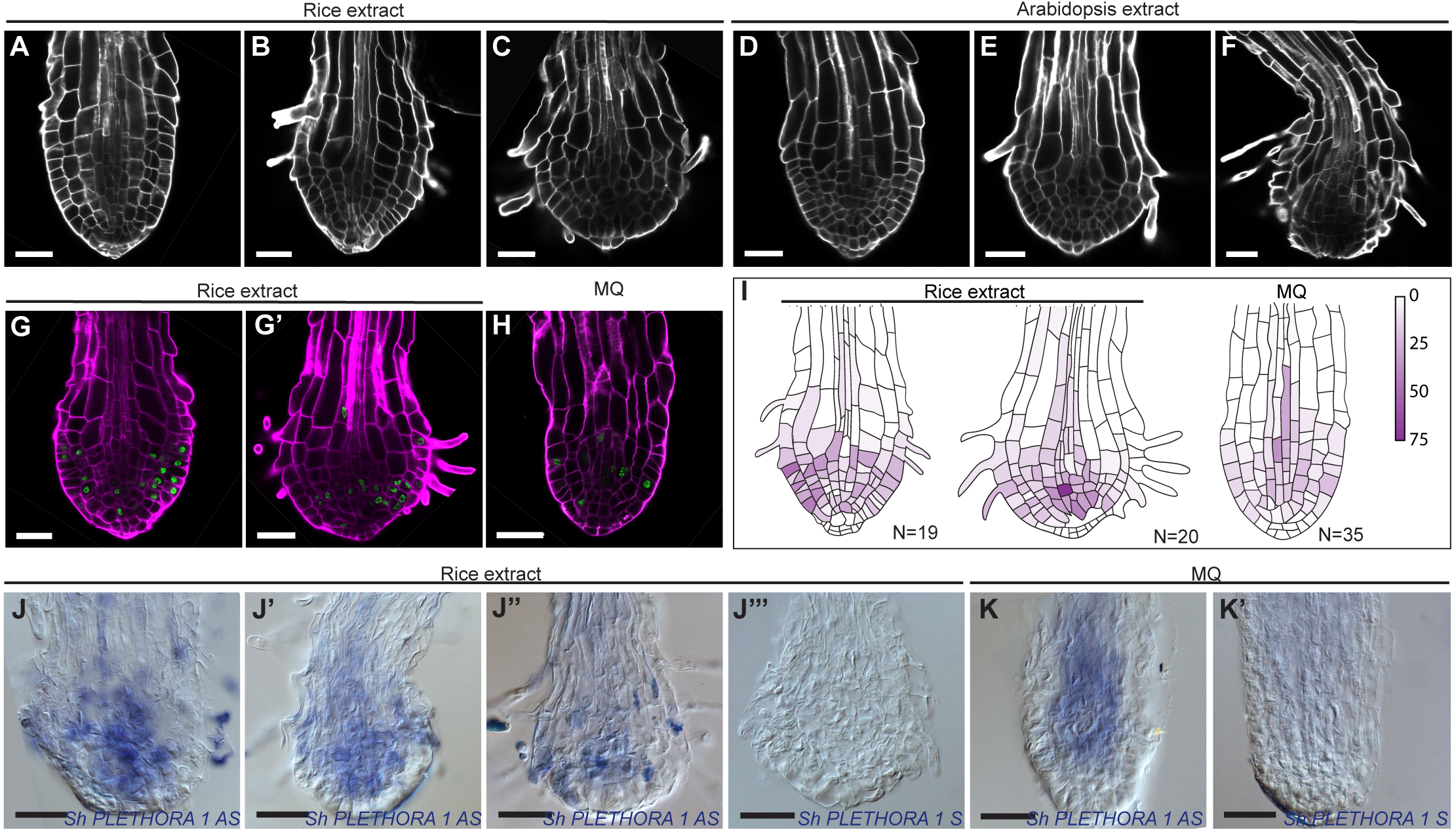
Prehaustorium of *Striga hermonthica* induced by host and non-host root extract treatment. A-F, Prehaustorium development after treatment with rice (A-C) and *Arabidopsis* extracts (D-F). Roots were treated after 24 h of germination and imaged after 4, 8, and 12 h. G-H, EdU staining showing cell division in the differentiating *S. hermonthica* roots. I, quantification of the cell division rate during prehaustorium formation, quantification of the cell division rate during root differentiation, purple bar indicates the percentage of chance of that cell may undergo cell division (EdU marked nucleus /N*100). J-K’, RNA *in situ* hybridization during prehaustorium formation in longitudinal sections using *ShPLETHORA1*. (J-K’), 24-h germinated seedlings were treated for 12 h with rice extract. Scale bars: 50 μm.

### Prehaustorium formation is correlated with changes in auxin and cytokinin levels

Because *PLT* expression is regulated by auxins (Mähönen et al., 2014), the decrease in *PLT1* levels in *Striga* roots during the transition to the prehaustorium stage prompted us to investigate whether an alteration in the auxin distribution pattern also contributes to meristem differentiation. In the *Arabidopsis* meristem, auxin maxima define cell type organization and dictate root zonation during root growth (Grieneisen et al., 2007; Dello loio et al., 2008; Sabatini et al., 1999). Root zonation is modulated by the antagonistic effect of auxins and cytokinins, with high levels of cytokinins promoting elongation and differentiation and high auxin levels inducing cell division (Moubayidin et al., 2010). Therefore, we sought to visualize auxin and cytokinin distribution during *Striga* prehaustorium formation using anti-indole-3-acetic acid (IAA) (Mravec et al., 2017) (Fig. **S6)** and anti-cytokinin antibodies (Agrisera, AS09 437). Using immunolocalization in paraffin-embedded sections of *S. hermonthica* roots grown in water or with rice exudates, we monitored IAA distribution during root growth (Fig. **4A-D’**). In both water and rice exudates, the anti-IAA antibodies showed a high signal at the pre-elongated stage (Fig. 4B,B’). This signal became restricted to only a few cells at the root tips prior to root differentiation (Fig. **4B;B’;D;D’**).

**Fig. 4:**
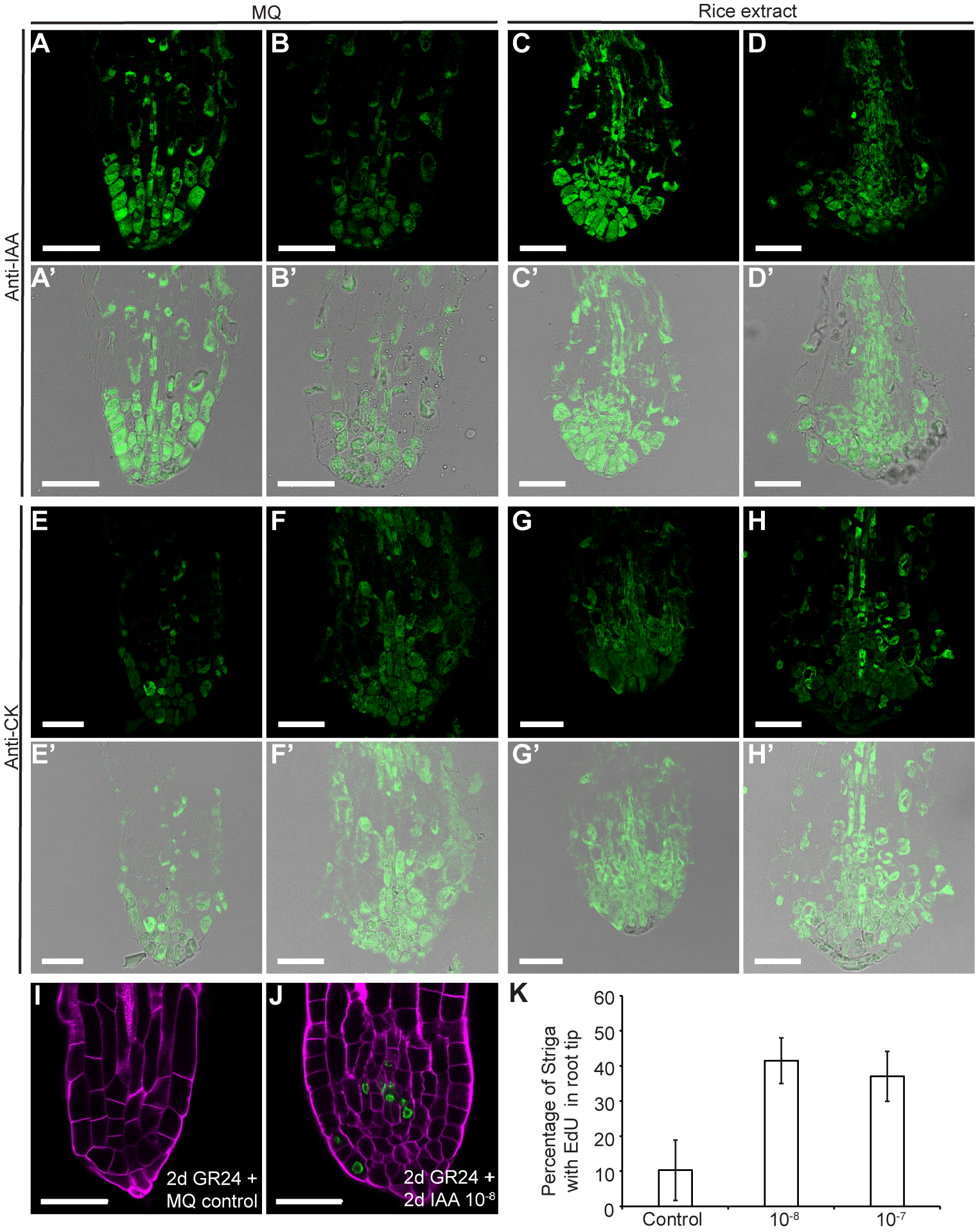
Auxin-cytokinin homeostasis control of *Striga hermonthica* meristem activity. A-D’, immunolocalization showing auxin distribution in *S. hermonthica* root sections using anti-IAA antibody, N = 30; E-H’, cytokinin accumulation detected using anti-cytokinin antibody. Seedlings are 24 (A;E) and 48 h (B;F) after GR24 treatment-induced germination and 12 h of rice extract treatment. N = 20; l-K, IAA application delayed meristem differentiation in the *S. hermonthica* radicle, as monitored by EdU staining. I and *J,* Confocal images of control (I) and IAA-treated roots (J). K, Percentage of cell division (EdU-stained cells) in *S. hermonthica* roots treated with different IAA concentrations. Columns indicate the means generated from three biological replicates; error bars represent SE. The number of *S. hermonthica* seedlings used were as follows for the control (N = 44; 53; 40); 10^-8^(N = 43; 45; 48); 10^-7^ (N = 93; 107; 95). Scale bars: 50 μm.

The anti-cytokinin antibodies exhibited an antagonistic pattern to auxins (Fig. 4E-H’). The anti-cytokinin signal was observed in a larger population of cells at the differentiation stage compared with that of anti-lAA antibodies (Fig. **4F;F’;H;H’**). The basic levels of auxins and cytokinins in the plant tissues were higher in roots treated with rice exudates than in those grown in water.

Low levels of auxin have been shown to stimulate root growth and delay root differentiation (Dastidar et al., 2019). Hence, we investigated whether the application of auxin at low concentrations prevents *Striga* meristem differentiation and delays haustorium formation. Because we did not observe striking differences between *S. hermonthica* roots grown in water and those grown in HIF and to assess the specificity of auxin, we used *S. hermonthica* grown in water. We incubated young *S. hermonthica* (48 h after induction with GR24) with different IAA concentrations and monitored root growth and cell division rates within the meristem using EdU (Fig. **S7;** Fig. **4I-K**). Three days after germination (dag), only 18% of untreated *S. hermonthica* showed active division in the meristem. This proportion was much higher following the application of low IAA concentrations for 1 day and was approximately 40% at 10^-8^, 42% at 10^-7^, and 32% at 10^-6^ M IAA. After treatment for 2 days (4 dag), only 6% of the control seedlings retained an active meristem, whereas treated *S. hermonthica* maintained their meristem with 39% at 10^-8^, 44% at 10^-7^, and 29% at 10^-6^ M IAA. At day 3 after auxin application (5 dag), we observed 17% at 10^-8^, 6% at 10^-7^, and 15% at 10^-6^ M IAA. IAA at a higher concentration, i.e., 10^-5^ M, inhibited cell division and elongation (Fig. **S7).** As *S. hermonthica* exhibited the highest rate of cell division on day 2 and at 10^-8^ and 10^-7^ M IAA, we focused on this time point and monitored cell division rates using 10^-8^ and 10^-7^ M IAA, which confirmed that auxin delayed *S. hermonthica* root differentiation (Fig. **4K**).

### A shift in PIN polarity precedes prehaustorium formation

In *Arabidopsis,* the PIN proteins, which serve as auxin efflux facilitators, play a major role in controlling the root meristem size by maintaining auxin maxima through establishing an auxin reflux loop in the meristem (Blilou et al., 2005). Furthermore, tissue-specific PIN polarity at the plasma membrane acts as a determinant for auxin flow directionality (Wisniewska et al., 2006). To determine whether the loss of meristematic activity during prehaustorium formation in *Striga* roots is also modulated by changes in PIN expression and polarity, we monitored the expression of PIN mRNA by *in situ* hybridization using *Striga PIN1* and *PIN2.* We also evaluated PIN1 and PIN2 polarity using *Arabidopsis-demed* PIN1 (Gälweiler et al., 1998) and PIN2 antibodies (Müller et al., 1998) derived from *Arabidopsis* in *S. hermonthica* roots grown in water or in the presence of rice extract.

We identified *Striga* PIN homologs using the recently published *Striga* genome sequence (Yoshida et al., 2019) (Fig. **S8).** RNA *in situ* hybridization using *ShPIN1-1* and *ShPIN1-2* probes, which target the mRNA of two *AtPIN1* homologs, showed that the two *Striga PIN1* orthologs have distinct expression patterns compared with *Arabidopsis. We* detected *ShPIN1-1* expression in the epidermis, cortex, and vasculature in the meristem (Fig. **5A-A”’**). *ShPIN1-2* mRNA expression could not be detected in the meristem, and we observed the signal above the meristem region (Fig. 5B-B”’). Next, we monitored PIN2 expression, and, unlike *Arabidopsis* PIN2, which is usually expressed in the cortex and epidermis, we detected high *ShPIN2* expression levels in the vasculature (Fig. **5C-C’”**).

**Fig. 5:**
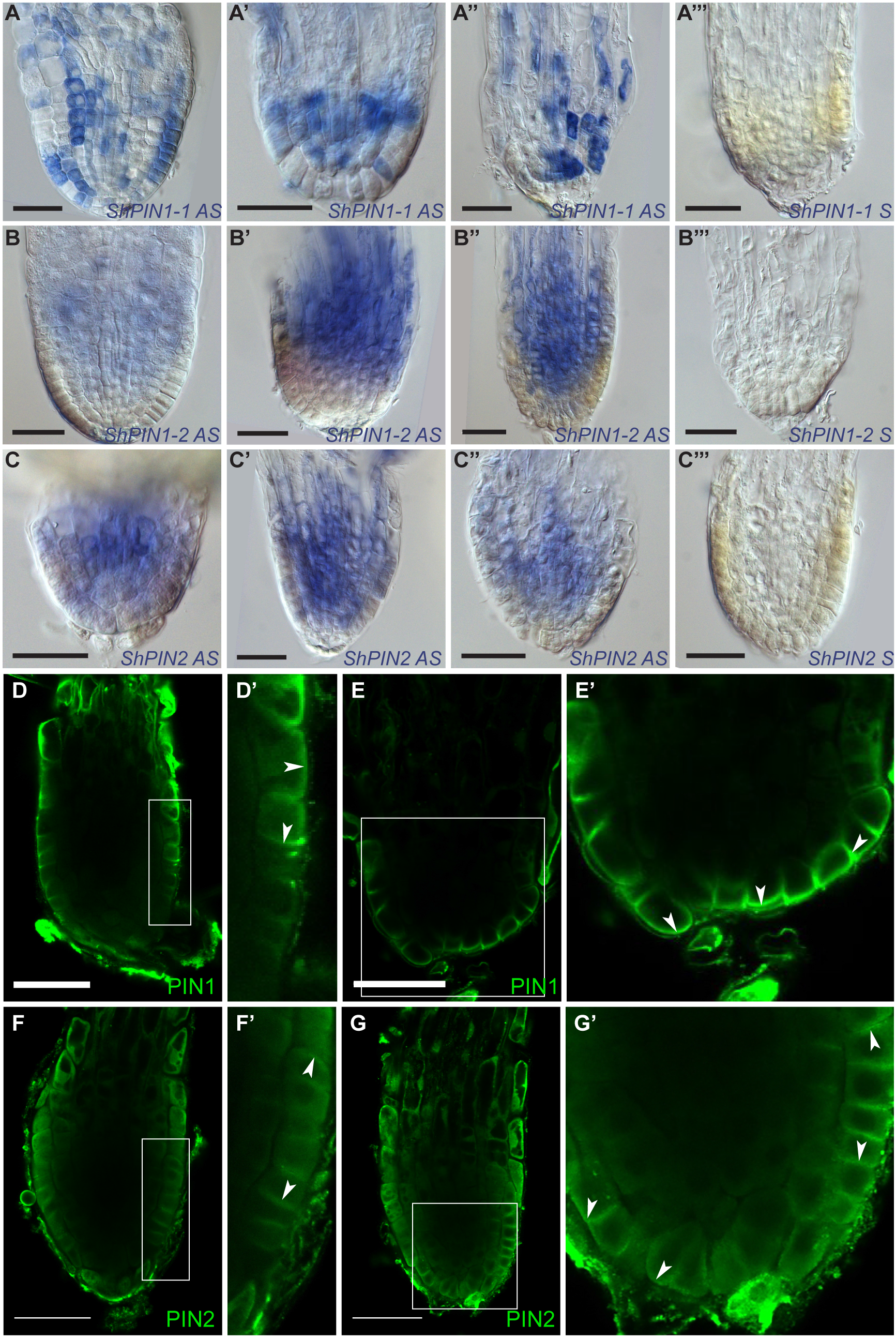
PIN protein expression and polarity in *Striga hermonthica* during root differentiation. A-C’”, RNA *in situ* hybridization of *S. hermonthica* root meristems at the young, pre-elongated, and elongated stages in longitudinal sections using *ShPIN1-1* (A-A’”), *ShPIN1-2* (B-B”’), and *ShPIN2* (C—C”’). A-A” and B-B” are the antisense probes and A’” and B”’ are the sense control probes. D-G’, Immunolocalization using *Arabidopsis* thaliana PIN1 and PIN2 antibodies showing (D-E’) basal PIN1 in the epidermis. D’ and E’ are insets from D and E, respectively. N = 30. F-G’, PIN2 localization showing epidermal apical, lateral, and basal polarity. Arrows indicate the shift in PIN polarity. F’ and G’ are insets from F and G, respectively. *S. hermonthica* seedlings were treated with GR24 for 24 or 48 h. N = 60. Scale bars: 50 μm.

In the *Arabidopsis* root meristem, PIN1 is localized in a polar manner at the basal end of the vasculature cells (Gälweiler et al., 1998). PIN2 is expressed in the cortex and epidermis and localizes in a polar manner at the apical side of cells in the epidermis and basally in the cortex (Blilou et al., 2005). To evaluate PIN polarity in *Striga* roots, we monitored the localization of *Arabidopsis* anti-PIN1 and anti-PIN2 antibodies in *S. hermonthica* using immunolocalization. In plants grown in water, we did not detect *Arabidopsis* PIN1 in the vasculature but rather in the epidermis with lateral and basal polarity at the early stage (Fig. **5D;D’**). At the elongated stage, PIN1 was predominantly located basally at the tip of the root from where the hairs emerge (Fig. 5E;E’). We found apical and basal PIN2 localization in the epidermis at the early stage (Fig. **5F;F’**). At the elongated stage, PIN2 was located basally and laterally outward from the epidermis and the cells located at the columella (Fig. **5G;G’**).

### Auxins are depleted from *Striga* roots during prehaustorium formation

From the elongated stage, PINs displayed a shift in polarity toward basal protein localization in cells located at the root cap position (Fig. 5). To predict the dynamics of auxin distribution in the *Striga* root during growth, we adapted an established mathematical model for *Arabidopsis* roots (Savina and Mironova, 2020) based on *Striga* root developmental layouts (Figs 1;2) and PIN1 and PIN2 expression patterns and polarity (Fig. 5). The model considers mutual correlations between auxin and PIN dynamics issued from positive and negative self-regulations of auxin transport (Supplementary information) (Fig. 5). We implemented the principle difference between *Striga* and *Arabidopsis* models in which PINs were able to localize on the outer membranes at the root cap position. With these localization patterns and in the absence of auxin reflux, we anticipated that auxin would be excreted from the roots to the environment.

Accordingly, we developed three two-dimensional cell layouts mimicking longitudinal root sections at the early young, elongated, and pre-elongated stages (Fig. S1, Fig. S9). The model parameters were adjusted to obtain auxin distribution, with an *Arabidopsis-*like auxin pattern having the maximum at the QC and high auxin levels in the root cap (Fig. 6A;D;G, Fig. S10 A-D). This was obtained when the negative regulation of PIN1/2 expression at high auxin levels and positive regulation at low levels were considered. Next, we entered the generated pattern at the young stage for PINs and auxin into the cell layout to the pre-elongated stage where cells elongated but without an increase in the cell division rate (Fig. S9) and continued the calculations without changing the model parameters or equations. As a result of auxin dilution caused by cell elongation, we observed both a decrease in the auxin maximum (Fig. 6B, Fig. S10) and a shift in PIN localization domains toward the root cap (Fig. 6E;H). The auxin levels in the QC reduced to those observed for the proximal meristem at the early stage, in which active cell division occurs (Fig. 6J;K). The reduction in auxin levels promotes cell division. Moreover, with significant PIN1 expression in cells at the root cap position (Fig. 6M), it begins excreting auxin into the environment and depleting auxin reserves from the root tip (Fig. 6L). Interestingly, we observed a robust correlation for the PIN1 and PIN2 expression domains between the simulation (Fig. 6D-I, Figs. S9;IO) and experimental results (Fig. 5). When the calculations were implemented for the elongated stage layout, we found high PIN1 and PIN2 expression in the cells located at the root cap position where auxin was excreted more efficiently from the *S. hermonthica* roots (Fig. 6, Fig. S9).

**Fig. 6:**
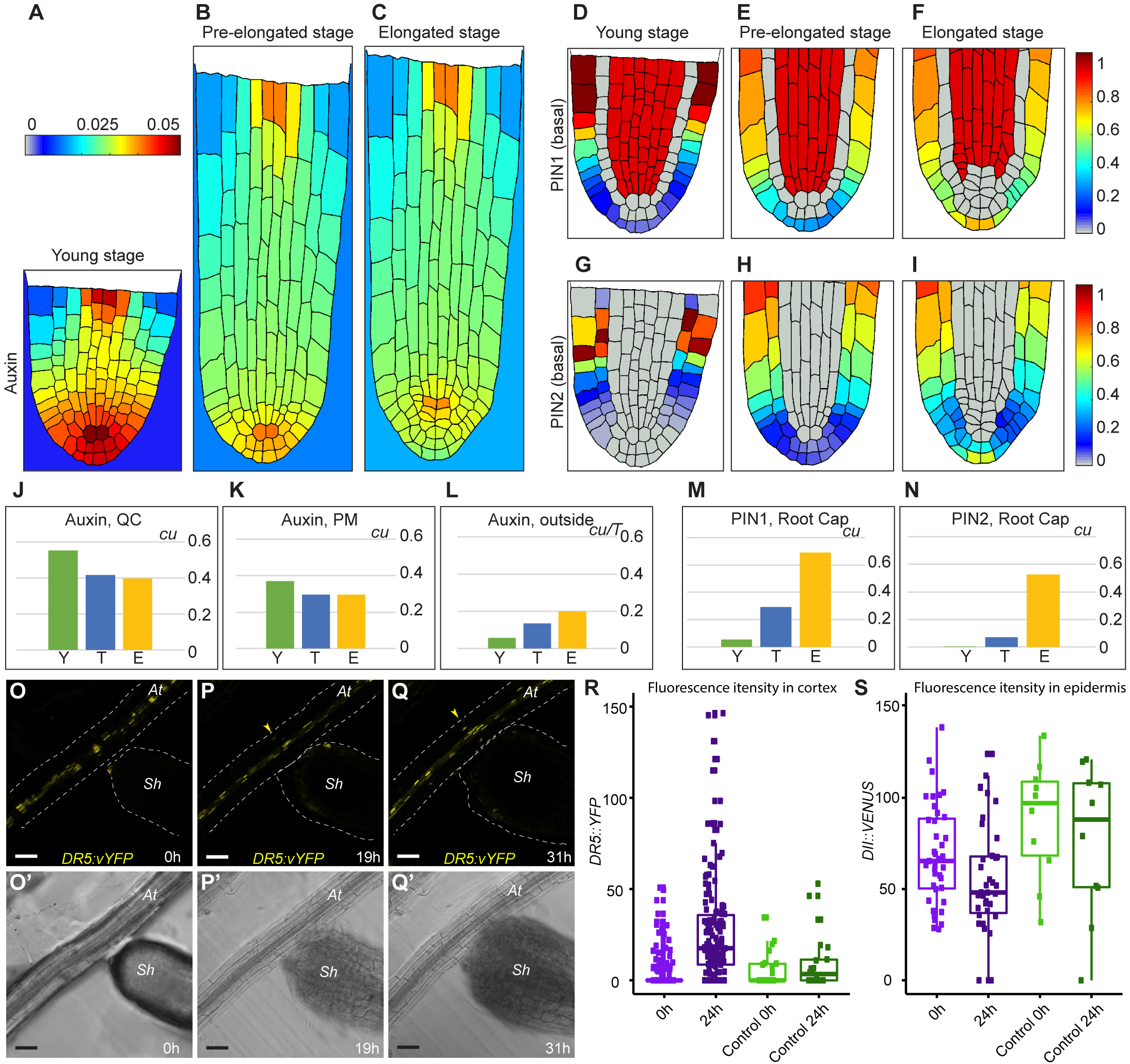
Mathematical model depicting auxin distribution in *Striga hermonthica* during root growth. (A–C), Quasi-steady state in auxin distribution inside and outside the *Striga* root at three consequent developmental layouts. (A,D,G) Young stage (Y), (B,E,H) pre-elongated stage (T), and (C,F,I) elongated stage (E). (D–F) PIN1 expression, (G–I) PIN2 expression with basal polarity. PIN2 with apical polarity is depicted in Fig. S9. (J–N) Comparison of the variable values between the stages. Auxin levels in the QC (J); in the proximal meristem (PM) (K); excreted from *S. hermonthica* roots (L) during calculation time *T*. m, PIN1 levels in the root cap; n, the level of basally polarized PIN2 in the root cap. cu, concentration units. O–Q’, time-lapse images of germinating *S. hermonthica* in contact with *Arabidopsis* roots containing *DR5::vYFP. S. hermonthica* were used 24 h after germination, N = 10. O-Q, Fluorescence YFP channel. O’-Q’ PMT transmission channel. Arrowheads indicate ectopic *DR5::vYFP* expression in P, Q, at *A. thaliana; sh, S. hermonthica.* Scale bars: 50 μm. r, the DR5 values are significantly different between contact 0–24 h (two-tailed t-test, p-value = 0.03794) but not significantly different between no contact 0–24 h (two-tailed t-test, p-value = 0.4947). s, the DII values are significantly different between contact 0–24 h (two-tailed t-test, p-value < 2.2e-16) but not significantly different between no contact 0–24 h (two-tailed t-test, p-value = 0.3095).

Our model indicates that *Striga* excretes auxin during root differentiation/haustorium formation. Because *Arabidopsis* root exudates induced haustorium formation as efficiently as rice exudates (Fig. 3D-F; Fig. S3), we used the *Arabidopsis* auxin response reporter lines *DR5::vYFP* (high auxin response) and *DII::VENUS* (low auxin response) to evaluate whether *Striga* induces changes in auxin levels or distribution when in contact with *Arabidopsis* roots (Fig. 6R;S). Using time-lapse imaging, we evaluated and quantified the fluorescence levels in both the epidermis and cortex of A. *thaliana* roots in contact with *S. hermonthica* from the young stage to the formation of prehaustoria (Fig. 6O-Q’). We observed ectopic accumulation of *DR5::vYFP* fluorescence and reduction of the *DII::VENUS* fluorescence signal in the root epidermis and cortex when the prehaustorium began to be formed and was physically attached to *the Arabidopsis* root (Fig. 6R;S, Movie/video 1). No expression in the cortex was observed when the *S. hermonthica* root did not differentiate or when it failed to reach *the Arabidopsis* root (Fig. S11, Videos 2, 3). These results are in agreement with those of our model and support the idea that auxin is depleted from the meristem prior to haustorium formation.

## Discussion

Root parasitic weeds undergo developmental programming to form their invasive organ, termed the haustorium, which emerges from the roots (Joel, 2013). The haustorium is essential for colonizing the host and deploys haustorial hairs by combining mechanical pressure and excreted cell wall-degrading enzymes (Perez-de-Luque, 2013; Reiss and Bailey, 1998). The germination of parasitic plants and the formation of haustoria are the most studied events in these invasive species because they form a putative point of attack to fight the invasion of crop plants (Baird and Riopel, 1983; Okonkwo and Raghavan, 1982; Kountche et al., 2016). A recent study has focused on the mechanisms underlying haustorium formation (Yoshida et al., 2016). Many studies have focused on either the transcriptional response (Ranjan et al., 2014; Yang et al., 2014; Yoshida et al., 2019) or the developmental characteristics of lateral haustoria (Ishida et al., 2016; Wakatake et al., 2020), which develop through dedifferentiation from mature cells in the cortex, resembling lateral root emergence or nodule formation during symbiosis with rhizobia (Yoshida et al., 2016). In this study, we assessed the developmental programs governing the formation of terminal haustoria in *S. hermonthica.* This parasite causes massive losses in cereal production and endangers food supply, particularly in sub-Saharan Africa (Pennisi, 2015). Following the sequencing of its genome, *S. hermonthica* can serve as a model plant for parasitic plants invading the roots of crop plants (Ranjan et al., 2014; Yang et al., 2014; Yoshida et al., 2019). In many root parasitic plants, including the holoparasitic *Orobanche* and other *Striga* species, the development of terminal haustoria is essential for root colonization. As these parasitic plants have small seeds and constrained nutrient availability, their survival relies entirely on developing a terminal haustorium that invades the host shortly after germination and exploits and hijacks its nutrient supply. Our results provide a mechanism that explains the switch from the vegetative to the parasitic state, leading to the formation of terminal haustoria after germination.

Our tissue anatomy analysis, consistent with those of a previous study (Hood et al., 1998), revealed that *S. hermonthica* embryonic roots, regardless of whether they are grown with or without HIF, have a meristem organization similar to that of *A. thaliana* (Fig. 1). Although we detected a root cap in *S. hermonthica* roots (Hood et al., 1998), we did not observe root cap sloughing during root growth, unlike that in other plant species (Fig. S1). In nonparasitic plants, the release of outer root cap cells into the soil decreases the frictional resistance (Bengough and McKenzie, 1997), but in *Striga,* this function might be required only for a short period of growth until reaching the host’s roots.

During the young stage, *S. hermonthica* roots possessed a meristem with active cell division, which is in accordance with the results of a previous study on *P. japonicum* roots (Ishida et al., 2016) During progression to the prehaustorium stage, we observed increased cell division in the stem cell niche area, including QC cells (Fig. 2). This is in contrast to the observations in *Arabidopsis,* in which QC cells undergo infrequent cell division. The observed increase in QC cell divisions might be a consequence of the accumulation of cytokinins in the distal meristem, as suggested previously (Zhang et al., 2013) (Fig. 3). Alternatively, these divisions might result from the induction of stress-induced signaling pathways. In *Arabidopsis,* elevated levels of the defense hormone jasmonic acid (JA) induce QC cell division (Zhou et al., 2019). Therefore, it is plausible that JA signaling contributes to the observed division in the *S. hermonthica* stem cell niche area, as supported by the enrichment in genes involved in response to wounding during haustorium formation (Ishida et al., 2016). The levels of reactive oxygen species (ROS) also play a role in modulating QC cell division (Lee, 2019; Yamada et al., 2020). Recent studies have reported an increase in peroxidase (precursor of ROS) activity during terminal and lateral haustoria formation (Wada et al., 2019; Yoshida et al., 2019). In most plants, the QC serves as a reservoir to replenish root stem cells during their growth; however, in *Striga,* QC signaling occurs transiently and might be required only for the establishment of the meristem at the young stage.

Although the progression to meristem differentiation in *S. hermonthica* roots was similar when the plants were grown in water or treated with HIFs, the cellular layout of the roots differed between these two conditions. When grown in water, *S. hermonthica* roots maintained their regular structure and displayed organized division planes (Fig. 1). When treated with HIFs, the cells divided in an unorganized manner (Fig. 3), which might have resulted from microtubule disorganization, as described during haustorium formation in the parasitic plant *Cuscuta* (Kaštier et al., 2018). With the complex composition of HIFs obtained from root exudates, many components have been shown to affect cell division, including terpenoids and flavonoids (Albrecht et al., 1999; Woo et al., 2005; Chaimovitsh et al., 2012). Upon progression to haustorium formation, we observed a reduction in the rate of cell division, followed by meristem differentiation (Figs. **2**;**3).** Our results corroborate those of transcriptomic studies showing downregulation of DNA metabolism genes during haustorium formation (Torres et al., 2005).

Our findings also revealed the importance of hormone homeostasis for prehaustorium formation (Fig. 4). In *Arabidopsis,* a balance between auxins and cytokinins dictates root zonation and an increase in cytokinin levels mediates meristem differentiation (Moubayidin et al., 2010). Additionally, the establishment of an auxin maximum through polar auxin transport dictates cell fate, cell division planes and rate, and cell polarity (Blilou et al., 2005; Sabatini et al., 1999). We showed that in *S. hermonthica* roots, differentiation at the stem cell region is correlated with a decrease in auxin levels at the root tip and an increase in cytokinin levels prior to haustorium formation (Fig. 4A-H’). We also demonstrated that the decrease in auxin levels at the root tip is correlated with a shift in PIN polarity (Fig. 5). In *Arabidopsis,* polar distribution of PIN proteins in the plasma membrane in a tissue-specific manner directs the auxin flux required for establishing the auxin maximum in the root meristem. Our investigation of *S. hermonthica* PIN proteins led to the two following findings: first, the PIN auxin efflux carrier detected by the *Arabidopsis* PIN2 antibody is not localized at the apical site of the epidermis in *S. hermonthica* but rather at the basal site of the cell. Second, the PIN transporter detected by the PIN1 antibody has distinct tissue localization compared with that in *Arabidopsis.* These differences suggest that *S. hermonthica* might use different polarity determinants in the epidermis. Our results are consistent with those of recent studies regarding the localization of PjPIN1 and PjPIN2 in *P. japonicum* during early haustorium formation (Wakatake et al., 2020). Both PjPIN1 and PjPIN2 accumulated at the basal end of cells toward the host cells (Wakatake et al., 2020). It will be essential to determine the localization of *Striga* PIN proteins to establish whether they have tissue type-specific polarity, as in *Arabidopsis.* However, such analysis will only be possible when a stable transformation protocol is available in Striga.

A study using a mesoscopic model in *Arabidopsis* established that a robust auxin maximum is generated by combining root topology, PIN protein distribution, and membrane permeability (Grieneisen et al., 2007). The concentration of PIN proteins depends largely on auxins, with PIN1 and PIN2 expression activated at a low auxin concentrations (Vieten et al., 2005). Our results using IAA antibodies indicated low auxin levels and high PIN1 and PIN2 levels at the epidermis. Our two-dimensional mathematical model recapitulated the decrease in auxin levels (Fig. 6B, Fig. S10) and a shift in PIN proteins toward the outer layers (Fig. 6E;G). Most importantly, with a decrease in auxin levels, the concentration of PIN1 increased in the outer layers, leading to auxin depletion in the root tip and meristem differentiation. Consistent with the decrease in auxin levels, exogenous IAA application delayed haustorium formation (Fig. 4). These results are in agreement with those of a previous study, in which exogenous auxin application poised the transition to the haustorium stage and cytokinin treatment induced haustorium formation (Keyes et al., 2001). Collectively, these results highlight the importance of asymmetric PIN protein distribution in the control of auxin flow and maintenance of the auxin maxima in the root meristem for directing tissue patterning and sustaining root growth.

Auxin excretion from *Striga* roots not only leads to *Striga* meristem differentiation and triggers haustorium formation, but may also contribute to host invasion. In *Arabidopsis,* auxins promote cell wall loosening by creating an acidic environment that stimulates cell wall remodeling enzymes (Majda and Robert, 2018). This strategy is also used by cyst and root-knot nematodes that can manipulate auxin signaling in their host plant to facilitate infection (Kyndt et al., 2016). It is plausible that the excreted auxins lead to an increase in auxin levels in the host cortex, which might contribute to cell wall degradation by activating host-specific cell wall-loosening proteins that facilitate host root invasion.

While our results provide a mechanistic understanding of terminal haustorium formation, transcriptome analysis during haustorium formation suggested an increase in auxin levels, indicated by the enrichment of genes involved in auxin signaling and transport, such as AUX/IAAs (Yoshida et al., 2019). It is plausible that the amount of auxins excreted from the roots during these stages in increased (Figs. **5**,**6**). We also observed an increase in cytokinin levels, which has been reported to induce haustorium formation and increase the aggressiveness of the parasitic plant *Phelipanche ramosa* (Goyet et al., 2019).

With severe *Striga* infestation in sub-Saharan Africa due to the large amount of seeds produced by each plant and their capacity to persist in the soil for almost a decade, one promising approach for reducing *Striga* invasion is suicidal germination, wherein synthetic stimulants are applied to the soil to induce germination in the absence of the host (Kountche et al., 2019). Preventing meristem differentiation by application of either auxin or a metabolite with a similar effect might be an alternative approach to suicidal germination by preventing haustorium formation in the presence of the host and thereby infestation by the parasite.

## Acknowledgments

This study was supported by King Abdulah University of Science and Technology (KAUST) baseline funding given to Ikram Blilou and by the Bill and Melinda Gates Foundation grant OPP1194472 given to Salim Al-Babili.

*We* thank Prof. Jiri Friml for generously providing AtPIN1 and AtPIN2 antibodies. We thank Dr. Wei Xu for the technical support during confocal imaging. Victoria Mirnova was supported by the Russian Science Foundation 18-74-10008. Savina Maria was supported by 0259-2019-0008-C-01.

## Authors’ contributions

The study was conceptualized by IB. The experimental designs were developed by Ikram Blilou, Ting Ting Xiao and Salim Al Babili. Ikram Blilou, Ting Ting Xiao and Gwendolyn K. Kirschner analyzed the *Striga* root anatomy and cell division rates. Ting Ting Xiao performed *in situ* hybridization and immunolocalization. Boubacar A. Kountche, Muhammad Jamil and Salim Al Babili contributed in the design of the auxin experiment, provided *Striga* seeds, helped with the sterilization and plant materials, and instructed Ting Ting Xiao and Gwendolyn K. Kirschner in handling *Striga* plants. Vinicius Lube performed *macro-Striga* images and edited movies from time-lapse imaging. Victoria Mirnova and Savina Maria performed mathematical modeling. All authors contributed to discussing the results of this manuscript.

## Conflict of interest

### Accessions

The genes used in this study have the following accession numbers: *ShPLT1* (StHeBC3_20835.1; GER38334); *ShHISTONE H4* (StHe2GB1_80290); *ShCYCLIN B1,3* (StHeBC3_3205); *ShPIN1-1* (StHe61GB1_24115; GER51912); and *ShPIN1-2* (StHe3G2B1_75771; GER32949).

---

**Fig. S1:** Cell division during *Striga hermonthica* development. A-D, cell wall staining using Renaissance stain. Purple arrowheads indicate the lateral root cap, N = 40. White arrowheads indicate divisions in the stem cell niche; blue arrowheads indicate divisions in the ground tissue; yellow arrowheads indicate root hairs. E-H, Lugol’s iodine stain indicates starch accumulation in *S. hermonthica* roots. Germination was induced with GR24 treatment and staining was performed 48 h after germination. N = 50. Scale bar: 50 μm.

**Fig. S2:** Schematic representation of the different stages of *Striga hermonthica* root development. Numbers inside the cells represent cells stained with EdU observed during the analysis respective to each stage; N represents the number of seedlings analyzed. Pareto chart plots of the distribution of the cell division ratio of each stage are shown in B.

**Fig. S3:** Phylogenetic trees of *Striga hermonthica, Arabidopsis,* and rice PLT proteins.

**Fig. S4:** *Striga hermonthica* prehaustorium formation with MQ water, *Arabidopsis,* and rice extract treatment. Yellow arrowheads indicate examples of structures considered to be prehaustoria. *S. hermonthica* treated with plant extract for 10 h after germination for 24 h. Scale bar: 200 μm.

**Fig. S5:** A-B, Schematic representation *of Striga hermonthica* prehaustorium formation induced with rice extract treatment. Numbers inside the cells represent cells stained with EdU; N represents the number of seedlings analyzed after 12 h of treatment with rice extract. C, Pareto chart plots of the distribution of the cell division ratio of each stage are shown in C. *S. hermonthica* treated with rice extract for 8 h (rice 1) and 10 h (rice 2) after 24 h of germination.

**Fig. S6:** Auxin treatment increases the IAA antibody signal in *Arabidopsis* root. Scale bar: 50 μm.

**Fig. S7:** EdU detection in *Striga hermonthica* root tips treated with different concentrations of IAA; number of *S. hermonthica* seedlings used are as follows for 3 dag: control (N = 54), 10^-8^ (N = 58), 10^-7^ (N = 62), 10^-6^ (N = 46), and 10^-5^ (N = 44); 4 dag: control (N = 44), 10^-8^ (N = 53), 10^-7^ (N = 40), 10^-6^ (N = 41), and 10^-5^ (N = 37); 5 dag: control (N = 39), 10^-8^ (N = 52), 10^-7^ (N = 29), 10^-6^ (N = 55), and 10^-5^ (N = 16); and 6 dag: control (N = 72); 10^-8^ (N = 69), 10^-7^ (N = 52), 10^-6^ (N = 42), and 10^-5^ (N = 63). dag, days after germination.

**Fig. S8:** Phylogenetic trees of *Striga hermonthica* and *Arabidopsis* PIN proteins.

**Fig. S9:** *Striga hermonthica* root structure layout used for constructing the mathematical model. A, young stage; B, pre-elongated stage; and C, elongated stage. All compartments are numbered. The cell layout in the pre-elongated stage was taken by transformation (elongation of the apical part) of the layout (A) without changes in the number of compartments. The cell wall types assigned to each cell are colored yellow (upward), red (downward), green (inward), and blue (outward).

**Fig. S10:** Auxin distribution pattern simulated in the mathematical model for the young stage (A-D), pre-elongated stage (E-H), and elongated stage (I-L). Quasi-steady state levels of auxins (a,e,i). PIN polarity with basal PIN1 (B,F,J) and PIN2 with basal (C,G,K) and apical (D,H,L) polarity in *Striga hermonthica* root tips. Only the cell layout changed between the stages. The model equations and parameters were the same for all calculations. Auxin concentration was calculated in the root and environment (per area) as PIN polarity allowed the excretion of auxins at the root tip.

**Fig. S11.** Contact of *Striga hermonthica* with *Arabidopsis* increases auxin levels in the epidermis and cortex. Three-dimensional time-lapse image of *S. hermonthica* reaching A. *thaliana* roots A-C’ and when failing to establish contact with the roots E-F’. N = 10. Scale bar: 50 μm

