## Supplementary Figure 1 for "The PLETHORA/PIN-FORMED/AUXIN network mediates terminal prehaustorium formation in the parasitic plant *Striga hermonthica*"

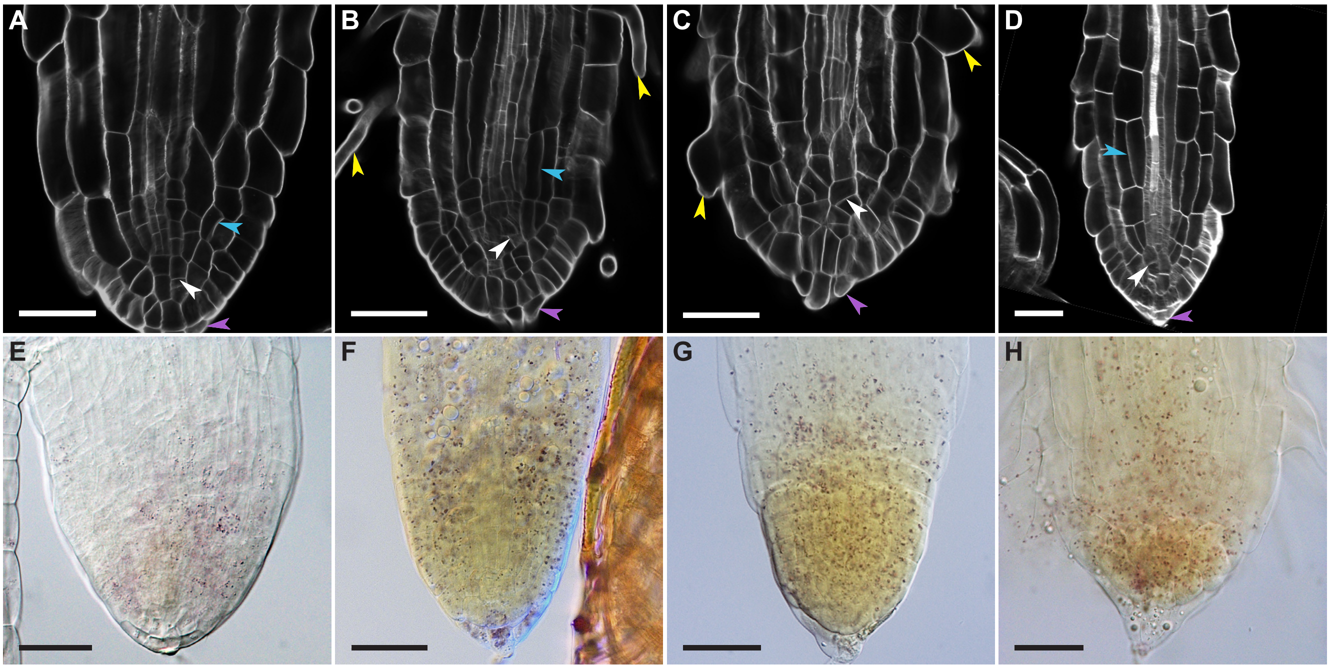


**Figure S1:** Ectopic cell divisions during Striga development. A-D, cell wall staining using Renaissance stain. Purple arrowheads mark the lateral root cap, N=40. White arrowheads point to divisions in the stem cell niche; blue arrowheads mark divisions in the ground tissue. Yellow arrowheads point to root hairs. E-H, Lugol stain marking starch accumulation in Striga roots, N=50. GR24 induced germination for 48h. Scale bars: 50µm.

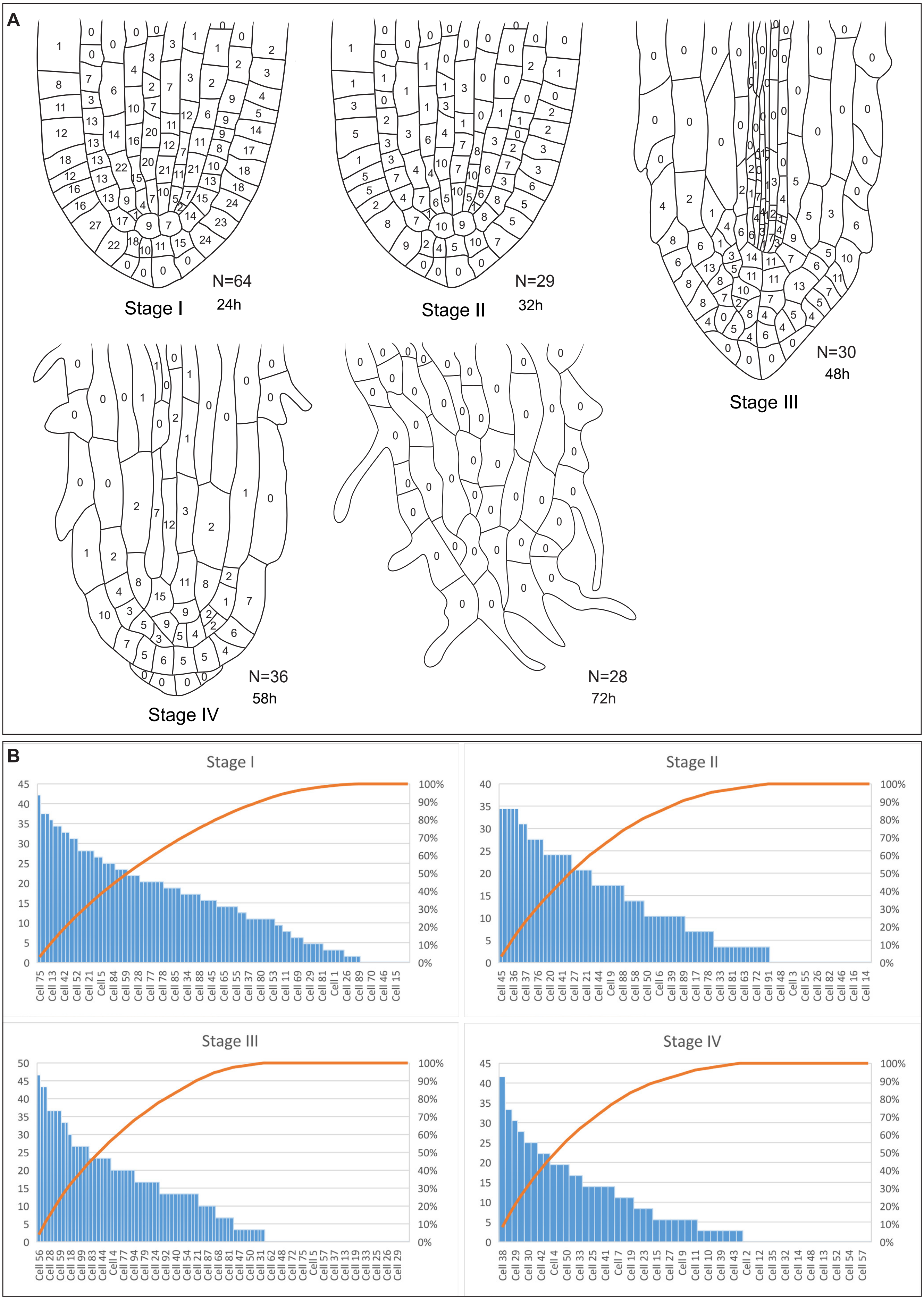


**Figure S2:** A, Schematic representation of different stages during Striga root development. Numbers inside the cells represent cells stained with EdU observed in the analyzed respective to each stage; N represents the number of seedlings analyzed. Pareto chart plots of the distribution of the cell division ratio of each stage are shown in B.

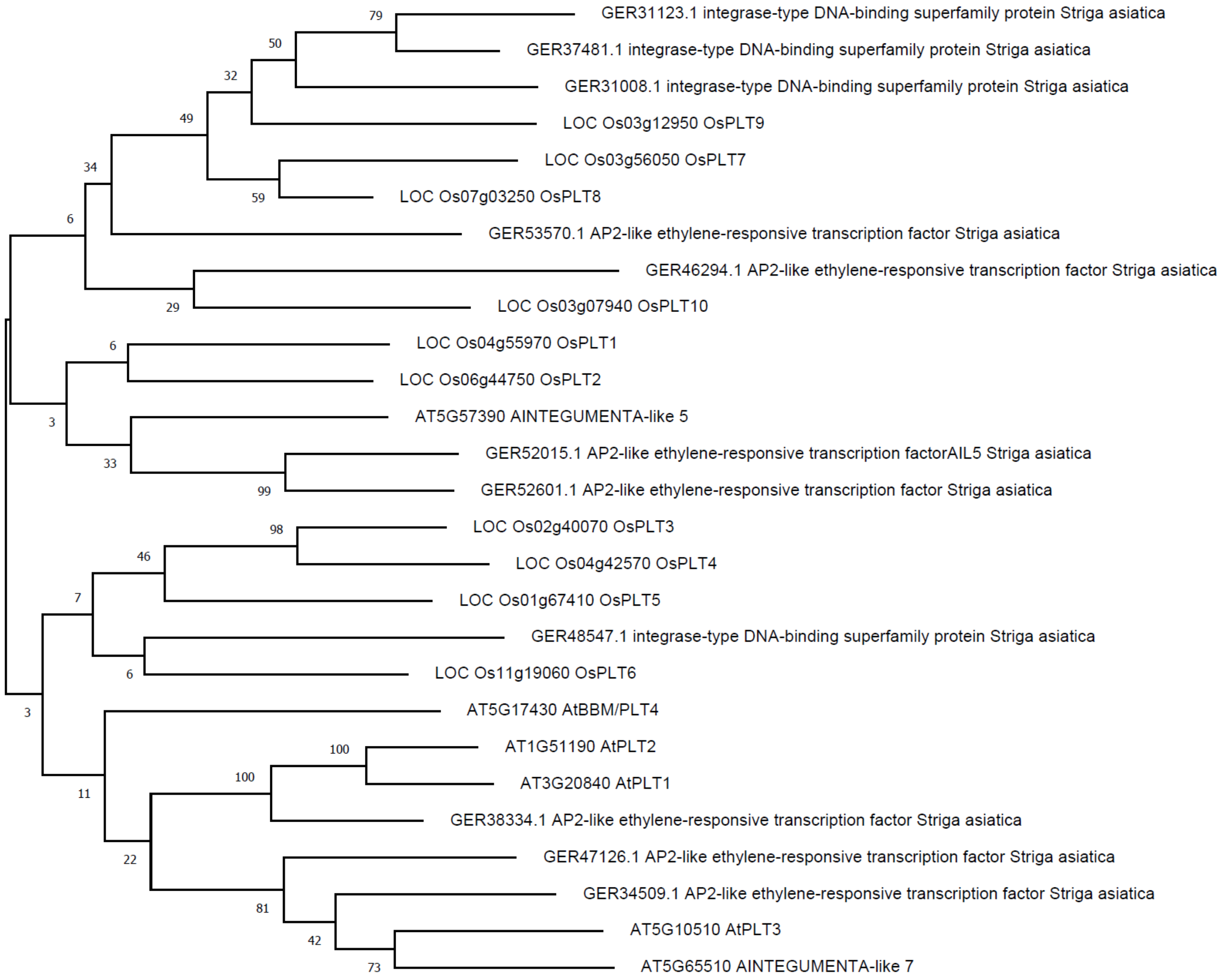


**Figure S3:** Phylogenetic trees of Striga, Arabidopsis, and rice PLT proteins.

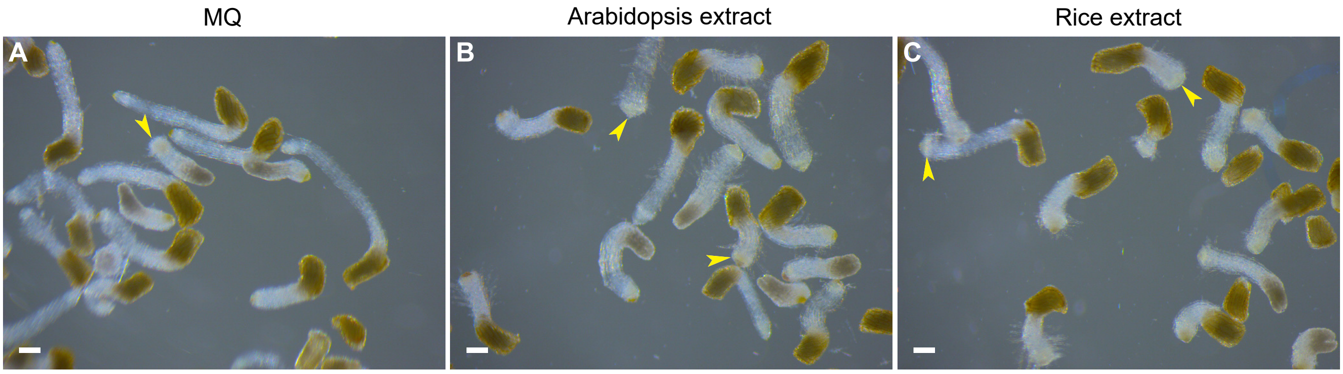


| Treatment | Total seedlings | Differentiated seedlings | Percentage of differentiated seedlings |
|---|---|---|---|
| MQ | 109 | 11 | 10.09174 |
| Arabidopsis | 145 | 86 | 59.31034 |
| Rice | 180 | 166 | 92.22222 |

**Figure S4:** Striga pre-haustorium formation with MQ, Arabidopsis and rice extracts treatment. Yellow arrowheads indicate the examples of structures considered as of pre-haustoria. Striga treated with plant extract for 10h after 24h germination. Scale bars: 200μm.

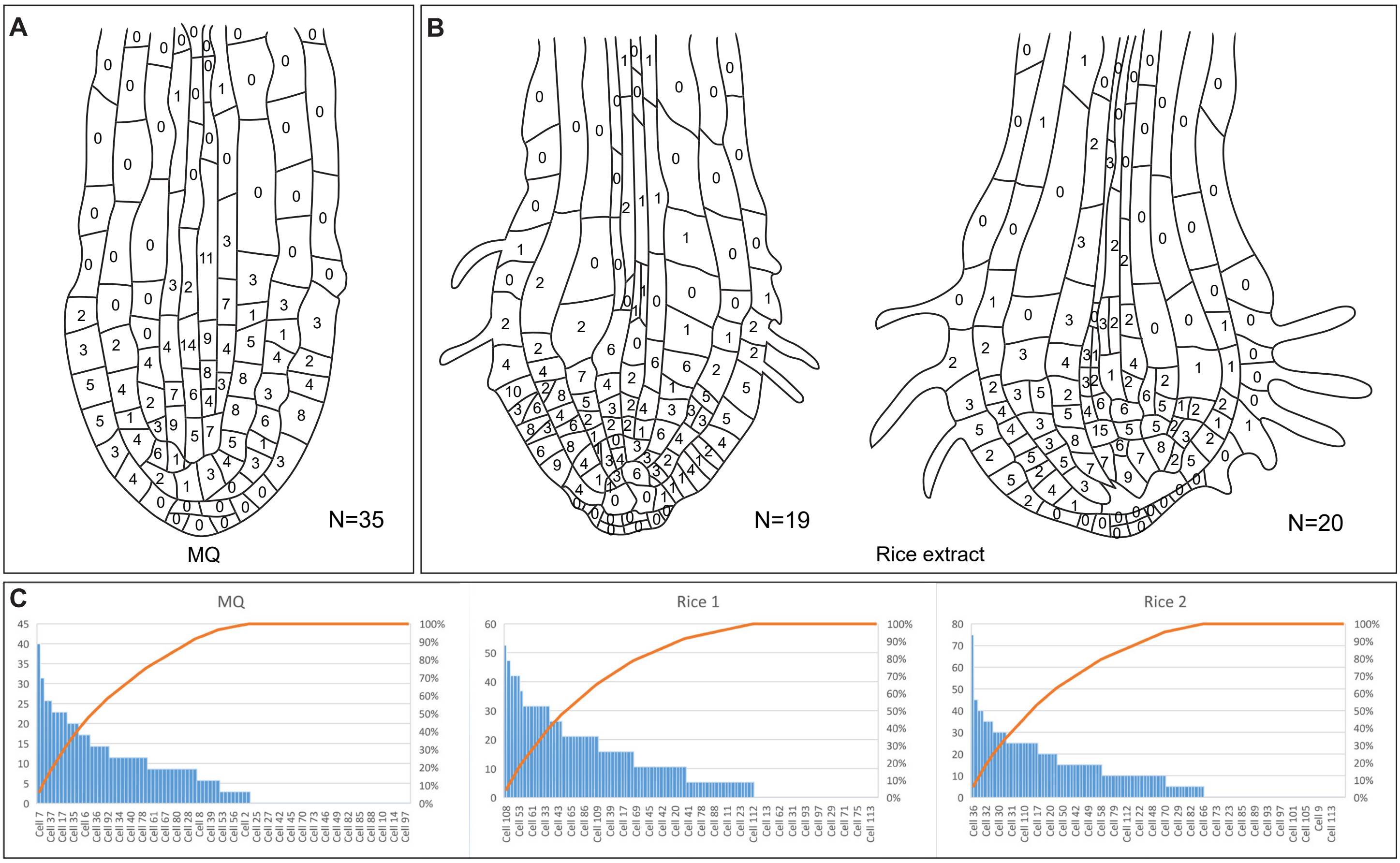


**Figure S5:** A-B, Schematic representation of Striga pre-haustorium formation induced with rice extract. Numbers inside the cells represent cells stained with EdU; N represents the number of seedlings analyzed; 12h rice extract treatment. C, Pareto chart plots of the distribution of the cell division ratio of each stage are shown in C.

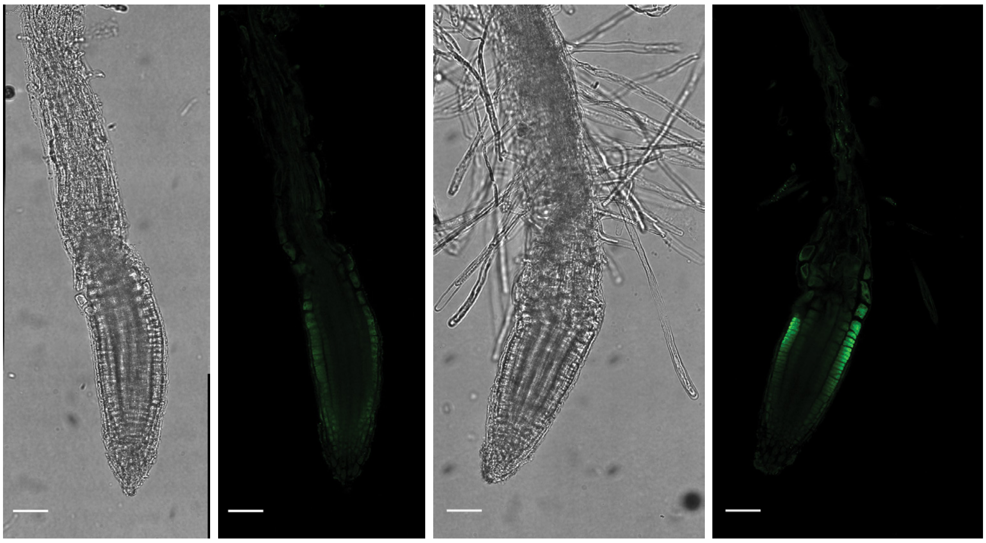

Control

100nM IAA treatment

**Figure S6:** Auxin antibody test on arabidopsis root.  Scale bars: 50μm.

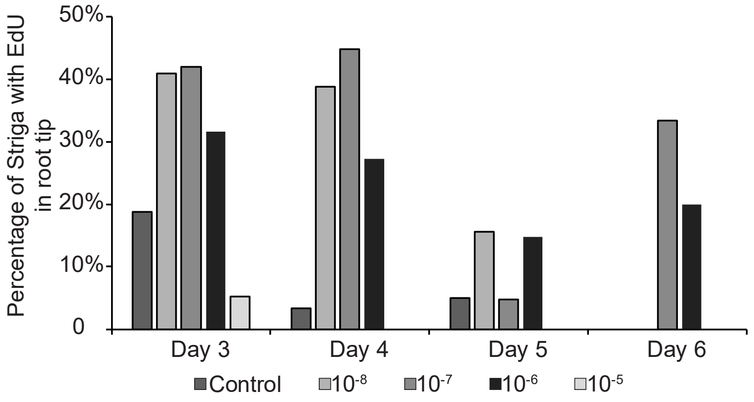


**Figure S7:** EdU detection in striga root tips after different concentration of exogenous IAA application; number of Striga seedlings used are as following for 3 dag: control (N=54); 10-8 (N=58); 10-7 (N=62); 10-6 (N=46) and 10-5 ( N=44); 4 dag, control (N=44); 10-8(N=53); 10-7 (N=40); 10-6 (N=41) and 10-5 ( N=37); 5 dag; control (N=39); 10-8 (N=52); 10-7 (N=29); 10-6 (N=55) and 10-5 ( N=16 ) and 6 dag control (N=72); 10-8 (N=69); 10-7 (N=52); 10-6 (N=42) and 10-5 (N=63); dag=days after germination.

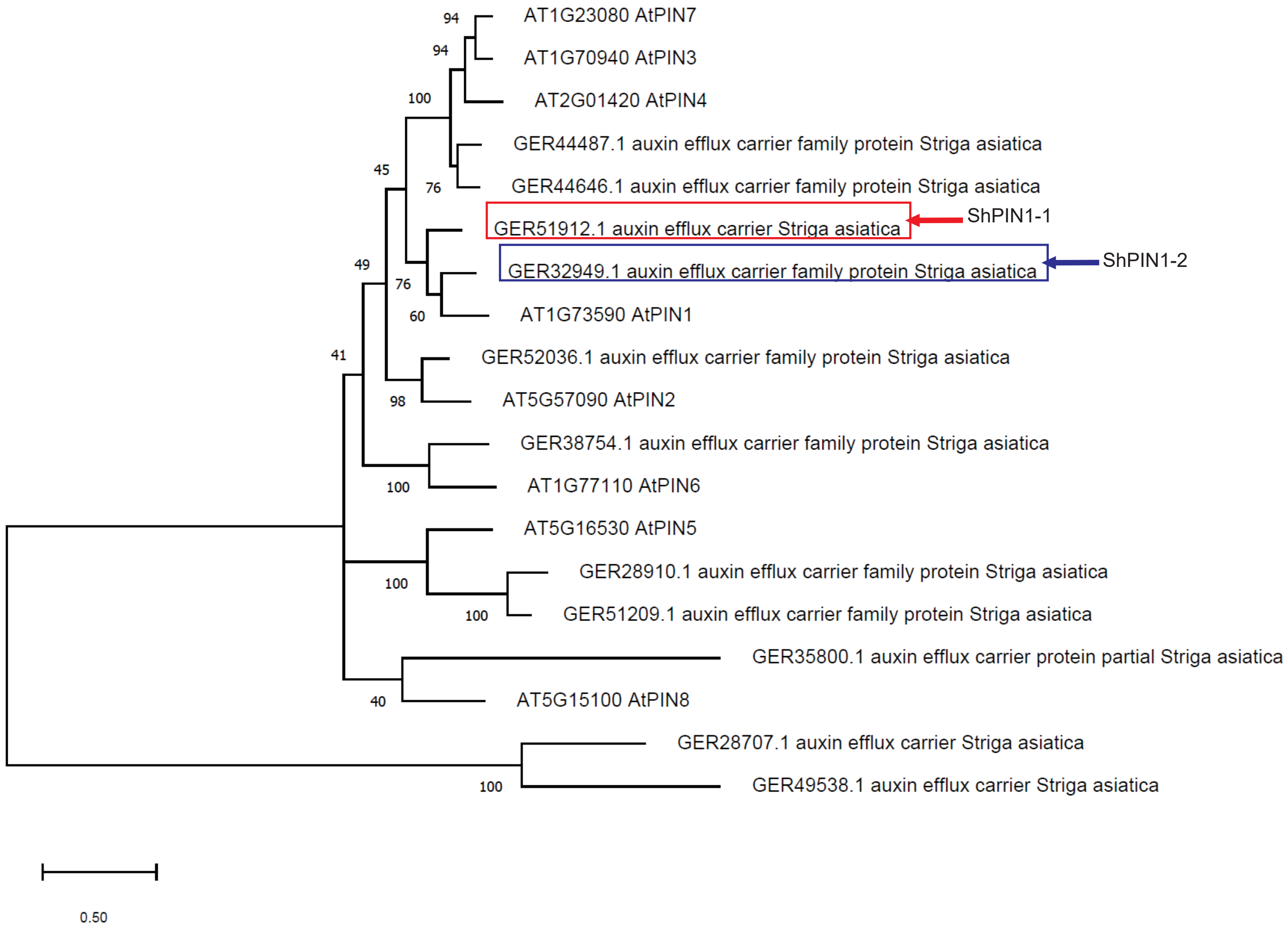


**Figure S8:** Phylogenetic trees of Striga and Arabidopsis PIN proteins

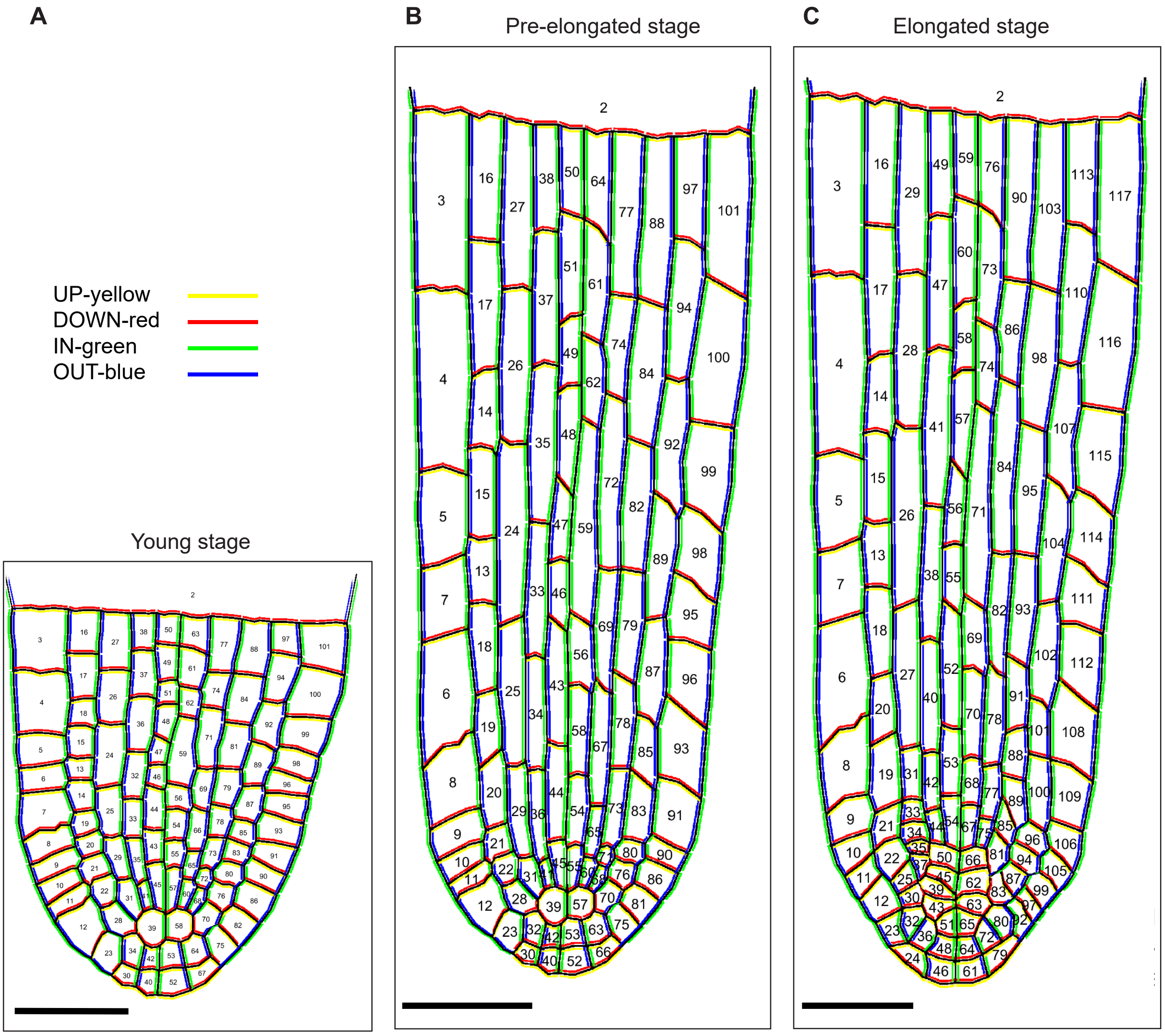


**Figure S9:** Striga root structure layout. A, young stage; B, pre-elongated stage; C, elongated stage. All compartments are numbered. The cell layout for the pre-elongated stage was taken by transformation (elongation of the apical part) of the layout (A) without changes in the number of compartments. The cell wall types assigned to each cell are colored by yellow (upward), red (downward), green (inward), and blue (outward).

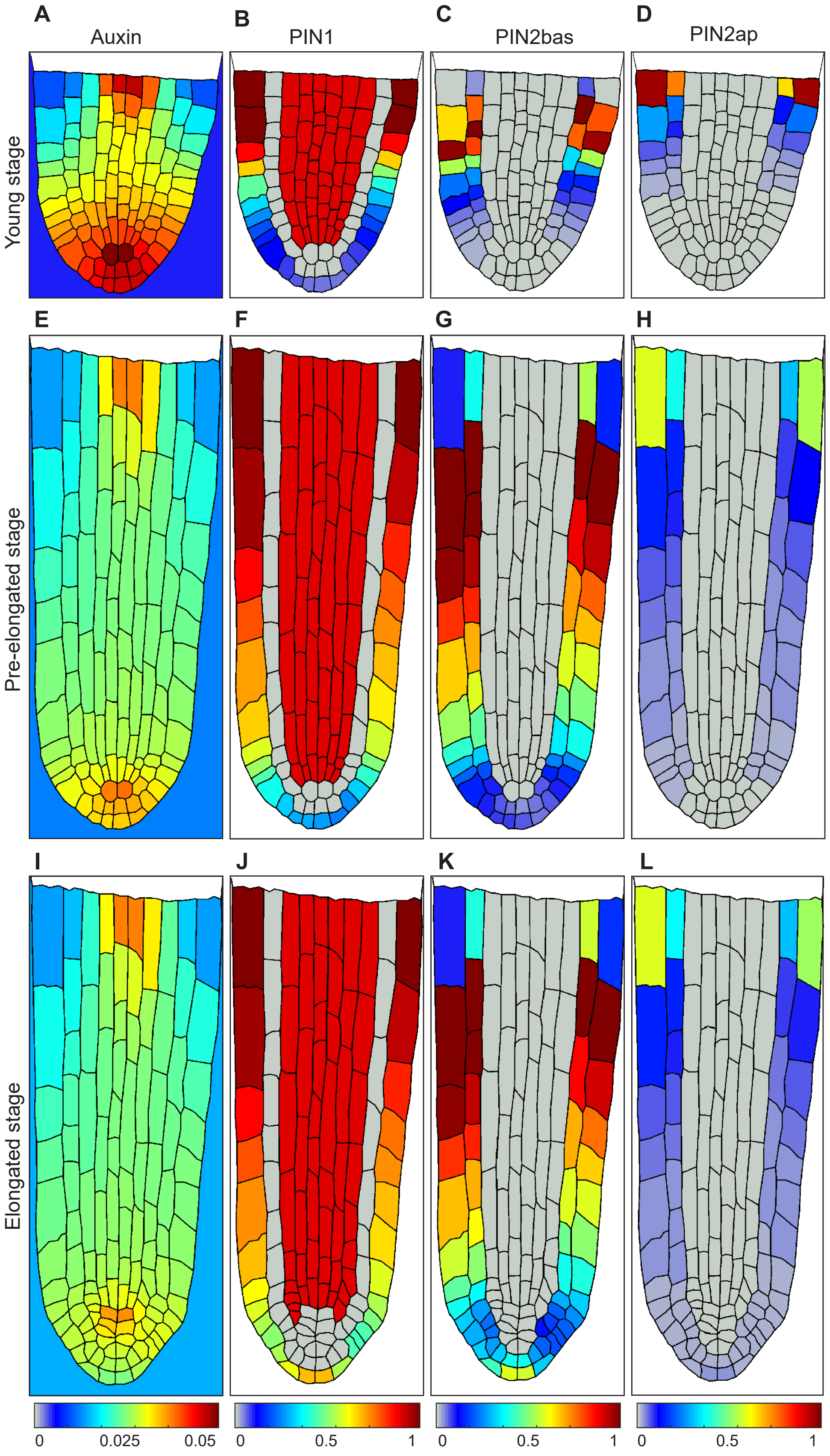


**Figure S10:** Auxin distribution simulated in the mathematical model for young stage (A-D), intermediate stage (E-H) and elongated stage (I-L). Quasi-steady-state levels of Auxin (A,E,I). PIN polarity with basal PIN1 (B,F,J) and PIN2 with basal (C,G,K) and apical (D,H,L) polarity in Striga root tip. Only cell layout changed in-between stages; the model equations and parameters were the same for all calculations. Auxin concentration was calculated in the root and the environment (per area), as PINs polarity allowed to excrete auxin at the root tip.

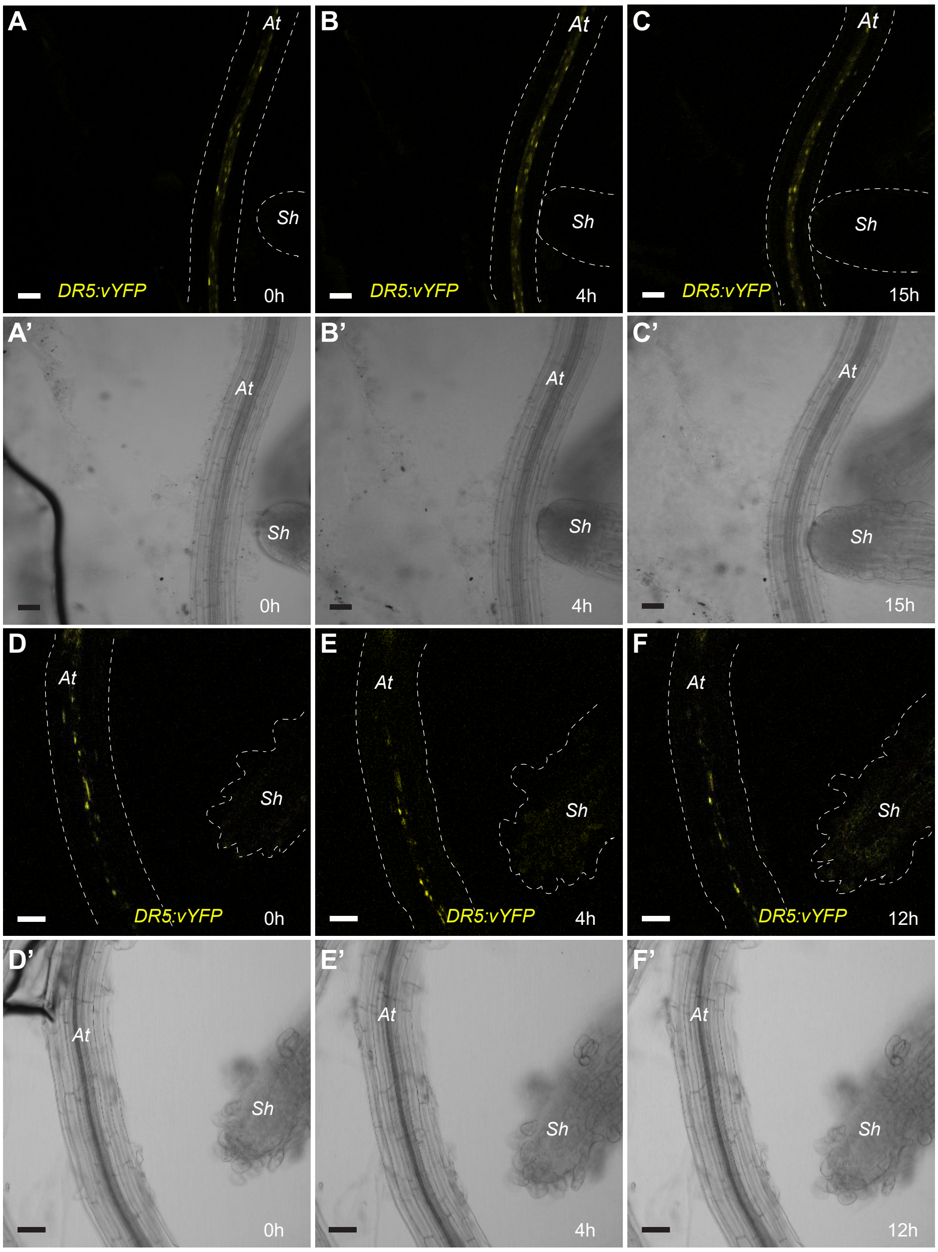


**Figure S11.** Striga contact with Arabidopsis is required to increase auxin. 3D time-lapse imaging of Striga reaching Arabidopsis roots A-C' and when failing to establish contact with the roots E-F'. N=10. Scale bars: 50µm.

**Supplementary Information**

**Mathematical modeling**

We constructed the mathematical model using PlantLayout pipeline [32] in MatLab ©. PlantLayout generates the structural model of a two-dimensional tissue anatomy, embeds the mathematical model in ODE, performs the numerical calculation, and visualizes the results using the structural model. The main details are described below.

1. Structural models

We constructed three structural models for *Striga hermonthica* root tips at three developmental stages Figure S5). The root tip layout for the young stage (A) was generated using a confocal image of the longitudinal section of the root tip. To generate the layout for the pre-elongated stage (B), we transformed the layout for the young stage by elongating the proximal meristem (until the initial vascular daughter cells), keeping the number of cells equal. The layout for the elongated stage (C) was a combination of the real root tip anatomy of a representative root at this stage and a generated proximal meristem from the intermediate stage. Such a design allowed the efficient transfer of the variable values between the layouts. PlantLayout calculated the volumes and relative positions of the cells as well as the length and orientation of the cell walls.

2. Mathematical model

The mathematical model considered the following processes: auxin degradation, diffusion (passive transport), PINFORMED (PIN) protein-mediated active transport, and auxin-dependent expression of PINs. There were four PINs in the model: PIN1in, PIN1out, PIN2ap, and PIN2bas. We assigned these to specific cell types and polarities. The vasculature and endodermis could express PIN1in with rootward (basal) polarity and fixed expression levels. The epidermis and Lateral root cap (LRC ) cells could express PIN1out with rootward (basal) polarity and auxin-dependent expression. The cortex, epidermis, columella, and LRC cells could express PIN2ap and PIN2bas with shootward (apical) and rootward (basal) polarities, respectively. The levels of shootward- and rootward-oriented PIN proteins in a cell dependent on auxin concentration.

The model in ODE is as follows:

$$\begin{cases} \frac{d[a]_i}{dt} = \left( k_\alpha * p_i + -K_{d,a}[a]_i + D \times \sum_{m \in N_i} \left( ([a]_m - [a]_i) l_{i,m} \right) + \sum_{x \in P} J_x(i) \right) \frac{1}{V_i} \\ \frac{d[PIN1in]_i}{dt} = 0 \\ \frac{d[PIN1out]_i}{dt} = \left( K_{s,PIN1out} \frac{\left( \frac{[a]_i}{q_{1,PIN1out}} \right)^{s_{PIN1out}}}{1 + \left( \frac{[a]_i}{q_{2,PIN1out}} \right)^{s_{PIN1out}}} - K_{d,PIN1out} \left( 1 + \left( \frac{[a]_i}{q_{3,PIN1out}} \right)^{h_{PIN1out}} \right) \times [PIN1out]_i \right) \frac{1}{V_i} \\ \frac{d[PIN2ap]_i}{dt} = \left( K_{s,PIN2ap} \frac{\left( \frac{[a]_i}{q_{1,PIN2ap}} \right)^{s_{PIN2ap}}}{1 + \left( \frac{[a]_i}{q_{2,PIN2ap}} \right)^{s_{PIN2ap}}} - K_{d,PIN2ap} \left( 1 + \left( \frac{[a]_i}{q_{3,PIN2ap}} \right)^{h_{PIN2ap}} \right) \times [PIN2ap]_i \right) \frac{1}{V_i} \\ \frac{d[PIN2bas]_i}{dt} = \left( K_{s,PIN2bas} \frac{\left( \frac{[a]_i}{q_{1,PIN2bas}} \right)^{s_{PIN2bas}}}{1 + \left( \frac{[a]_i}{q_{2,PIN2}} \right)^{s_{PIN2bas}}} - K_{d,PIN2bas} \left( 1 + \left( \frac{[a]_i}{q_{3,PIN2bas}} \right)^{h_{PIN2bas}} \right) \times [PIN2bas]_i \right) \frac{1}{V_i} \end{cases},$$

where $k_\alpha$ is the intensity of auxin flow from the elongation zone to the meristem; $p_i$ is the portion of auxin flow from the elongation zone per cell *i*; $K_{d,a}$ is the auxin degradation rate constant; *D* is the auxin diffusion rate constant; $N_i$ is the set of neighboring cells for cell *i*; $J_x(i)$ represents the auxin flow mediated by PINs, where *x* denotes PIN1in, PIN1out, PIN2ap, and PIN2bas; $K_{s,x}$ is the synthesis rate constant; $q_{1,x}$ is the activation threshold for auxin-dependent PIN protein synthesis; $q_{2,x}$ is the saturation threshold of auxin-dependent PIN protein synthesis; $s_x$ is Hill's coefficient, which determines the efficiency of PIN protein synthesis in response to changes in intracellular auxin concentration; $K_{d,x}$ is the degradation rate constant; $q_{3,x}$ is the threshold of auxin-dependent PIN protein degradation; $h_x$ is the coefficient that defines non-linearity of auxin-regulated PIN protein degradation; $V_i$ is the volume of cell *i* (in the two-dimensional case, the section area); and $l_{i,m}$ is the length of the cell wall between cells *i* and *m*.

The main source of auxins in the cell layout was the auxin flow from the elongation zone. It was assigned for the upper vasculature cells only, as follows (neighboring the index 2 space, Extended Figure 5):

$$p_i = \frac{l_{i,2}}{\sum_j l_{j,2}} \quad (2)$$

where $l_{i,2}$ is the length of the cell wall between cells *i* and 2, and $\sum_j l_{j,2}$ is the sum length of the cell walls of the stele and endodermis bordering the index 2 space.

The active auxin flows mediated by PINs were determined as follows:

$$J_x(i) = \sum_{m \in N_i} \left( k^x_{m,i} [a]_m [x]_m - k^x_{i,m} [a]_i [x]_i \right), \quad (3)$$

where $N_i$ is the set of neighboring cells for cell *i*; $k^x_{m,i}$ is the constant reflecting the portion of protein *x* located on the membrane of cell *m* neighboring cell *i* from the total protein *x* in cell *m*; and $k^x_{i,m}$ is the constant reflecting the portion of protein *x* located on the membrane of cell *i* neighboring cell *m* from the total protein *x* in cell *I*, *x*∈{ *PIN1in, PIN1out, PIN2ap, PIN2bas* }.

Parameters $V_i$, $l_{i,j}$, $k_{i,j}^{PIN1}$, $k_{i,j}^{PIN2}$, $k_{i,j}^{PIN3}$, $k_{i,j}^{PIN4}$, $k_{i,j}^{PIN7}$, and $p_i$ were defined by PlantLayout in the structural model (see above).

Boundary conditions: A specific feature of *S. hermonthica* is the ability of PINs to localize on the outer membrane of the root cap. Therefore, for LRC cells, we defined a specific boundary condition with no passive transportation and only active auxin transport depending on PIN1out and PIN2bas expression levels as follows:

$$J_x(i) = -k_{i,1}^{x}[a]_i[x]_i,$$

where x is PIN1out and PIN2bas, *i* is the LRC cell, and $k_{i,1}^{x}$ is the constant reflecting the portion of protein *x* located on the outer membrane of cell *i* neighboring the environment. There was no auxin exchange between the epidermis and the environment. The auxin level in the environment (compartment 1) is also calculated in the model as follows:

$$\frac{d[a]_1}{dt} = \left(\sum_{\substack{x\in P_1 \\ m\in N_1}} \left(k_{m,1}^{x}[a]_m[x]_m\right) - K_{d,a}[a]_1\right)\frac{1}{V_1},$$ where $P_1$=*{PIN1out, PIN2bas}*

3. Model calculation

First, we adjusted the model parameters for the young stage layout, such as auxin maximum located in the QC and PIN expression patterns corresponding to the experimental data. In this calculation, initial data were set to $[a]_i(0) = 0, [x]_i(0) = 0$ for *x* = PIN1out, PIN2ap, and PIN2bas. PIN1in(0) = 1. The parameter settings are presented in Table S1. The adjusted parameters (Table S1) were used without changes to model auxin distribution on the intermediate and elongated stages. The only changes made were to (1) the layout and (2) initial data. We used $[a]_i(T), [x]_i(T)$ concentration values taken from the quasi-steady state solutions at the previous stage as the initial data. *T* is the time of calculation to obtain the quasi-steady state. “Quasi” is due to the model being “open” with auxin leaking from the LRC.

**Table S1. Parameter settings used in the mathematical model**

| Parameter | Symbol | Units | Value |
|---|---|---|---|
| Intensity of auxin flow from the upper root part | $k_{\alpha}$ | *mu/tu* | 2,0 |
| Degradation rate constant for PIN1out | $K_{d,PIN1out}$ | *vu/tu* | 1000 |
| Degradation rate constant for PIN2ap | $K_{d,PIN2ap}$ | *vu/tu* | 1000 |
| Degradation rate constant for PIN2bas | $K_{d,PIN2bas}$ | *vu/tu* | 1000 |
| Degradation rate constant for auxin | $K_{d,a}$ | *vu/tu* | 0,005 |
| Synthesis rate constant for PIN1out | $K_{s,PIN1out}$ | *mu/tu* | 1000 |
| Synthesis rate constant for PIN2ap | $K_{s,PIN2ap}$ | *mu/tu* | 1000 |
| Synthesis rate constant for PIN2bas | $K_{s,PIN2bas}$ | *mu/tu* | 1000 |
| Diffusion transport rate constant | $D$ | *lu/tu* | 0,8 |
| Activation thresholds of auxin-dependent synthesis of PIN1out | $q_{1,PIN1out}$ | *1/cu* | 0,003 |
| Saturation thresholds for auxin-dependent synthesis of PIN1out | $q_{2,PIN1out}$ | *1/cu* | 0,0033 |
| Thresholds of auxin-dependent degradation of PIN1out | $q_{3,PIN1out}$ | *1/cu* | 0,03 |
| Hill coefficient that determines the rate of PIN1out synthesis in response to changes in intracellular auxin concentration | $s_{PIN1out}$ | *dl* | 2 |
| Coefficient that defines non-linearity of auxin-regulated PIN1out degradation | $h_{PIN1out}$ | *dl* | 6 |
| Parameters of auxin-dependent synthesis and degradation for PIN2ap | $q_{1,PIN2ap}$ | *1/cu* | 0,0015 |
| | $q_{2,PIN2ap}$ | *1/cu* | 0,00165 |
| | $q_{3,PIN2ap}$ | *1/cu* | 0,015 |
| | $s_{PIN2ap}$ | *dl* | 2 |
| | $h_{PIN2ap}$ | *dl* | 6 |
| Parameters of auxin-dependent synthesis and degradation for PIN2bas | $q_{1,PIN2bas}$ | *1/cu* | 0,0175 |
| | $q_{2,PIN2bas}$ | *1/cu* | 0,018 |
| | $q_{3,PIN2bas}$ | *1/cu* | 0,027 |

| | | | |
|---|---|---|---|
| | $s_{PIN2bas}$ | *dl* | 12 |
| | $h_{PIN2bas}$ | *dl* | 10 |

*mu*, mass units; *cu*, concentration units; *tu*, time units; *vu*, volume units; *lu*, length units; *dl*, dimensionless parameter.

**Table S1. Parameter settings used in the mathematical model**

| Parameter | Symbol | Units | Value |
|---|---|---|---|
| Intensity of auxin flow from the upper root part | $k_{\alpha}$ | *mu/tu* | 2,0 |
| Degradation rate constant for PIN1 | $K_{d,PIN1}$ | *vu/tu* | $1,0*10^{3}$ |
| Degradation rate constant for PIN2ap | $K_{d,PIN2ap}$ | *vu/tu* | $1,0*10^{3}$ |
| Degradation rate constant for PIN2bas | $K_{d,PIN2bas}$ | *vu/tu* | $1,0*10^{3}$ |
| Degradation rate constant for auxin | $K_{d,a}$ | *vu/tu* | $5,0*10^{-3}$ |
| Synthesis rate constant for PIN1 | $K_{s,PIN1}$ | *mu/tu* | $1,0*10^{3}$ |
| Synthesis rate constant for PIN2ap | $K_{s,PIN2ap}$ | *mu/tu* | $1,0*10^{3}$ |
| Synthesis rate constant for PIN2bas | $K_{s,PIN2bas}$ | *mu/tu* | $1,0*10^{3}$ |
| Diffusion transport rate constant | $D$ | *lu/tu* | $8,0*10^{-1}$ |
| Activation thresholds of auxin-dependent synthesis of PIN1 | $q_{1,PIN1}$ | *1/cu* | $3,0*10^{-3}$ |
| Saturation thresholds for auxin-dependent synthesis of PIN1 | $q_{2,PIN1}$ | *1/cu* | $3,3*10^{-3}$ |
| Thresholds of auxin-dependent degradation of PIN1 | $q_{3,PIN1}$ | *1/cu* | $3,0*10^{-2}$ |
| Hill coefficient that determines the rate of PIN1 synthesis in response to changes in intracellular auxin concentration | $s_{PIN1}$ | *dl* | 2,0 |
| Coefficient that defines non-linearity of auxin-regulated PIN1 degradation | $h_{PIN1}$ | *dl* | 6,0 |
| Parameters of auxin-dependent synthesis and degradation for PIN2ap | $q_{1,PIN2ap}$ | *1/cu* | $1,5*10^{-3}$ |
| | $q_{2,PIN2ap}$ | *1/cu* | $1,65*10^{-3}$ |
| | $q_{3,PIN2ap}$ | *1/cu* | $1,5*10^{-2}$ |
| | $s_{PIN2ap}$ | *dl* | 2,0 |
| | $h_{PIN2ap}$ | *dl* | 6,0 |
| Parameters of auxin-dependent synthesis and degradation for PIN2bas | $q_{1,PIN2bas}$ | *1/cu* | $1,75*10^{-2}$ |
| | $q_{2,PIN2bas}$ | *1/cu* | $1,8*10^{-2}$ |
| | $q_{3,PIN2bas}$ | *1/cu* | $2,7*10^{-2}$ |
| | $s_{PIN2bas}$ | *dl* | 12,0 |

| | $h_{PIN2bas}$ | *dl* | 10,0 |
|---|---|---|---|

*mu*, mass units; *cu*, concentration units; *tu*, time units; *vu*, volume units; *lu*, length units; *dl*, dimensionless parameter.

**Table S2: Primers used in this study**

| **Gene** | Striga identifier | Striga identifier | Primer forward | Primer reverse |
|---|---|---|---|---|
| *ShPLT1* | StHeBC3_20835.1 | GER38334 | ccgtgctgaatgttcccaag | gagacgagcgccagcatcac |
| *ShHISTONE H4* | StHe2GB1_80290 | | gatcagctactgttagc | ccggcgatccgtcgcctggc |
| *ShCYCLIN B1,3* | StHeBC3_3205 | | gggtcgtgtaattcatca | caatgacctggcggctgtcg |
| *ShPIN1-1* | StHe61GB1_24115 | GER51912 | ccaccaacgatccctacacc | cgggagagaacataccgttg |
| *ShPIN1-2* | StHe3G2B1_75771 | GER32949 | ccatcatggagtagaagtcgg | gaccatcaccctcttctc |
